# Energetics of stochastic BCM type synaptic plasticity and storing of accurate information

**DOI:** 10.1101/2020.01.28.922948

**Authors:** Jan Karbowski

## Abstract

Excitatory synaptic signaling in cortical circuits is thought to be metabolically expensive. Two fundamental brain functions, learning and memory, are associated with long-term synaptic plasticity, but we know very little about energetics of these slow biophysical processes. This study investigates the energy requirement of information storing in plastic synapses for an extended version of BCM plasticity with a decay term, stochastic noise, and nonlinear dependence of neuron’s firing rate on synaptic current (adaptation). It is shown that synaptic weights in this model exhibit bistability. In order to analyze the system analytically, it is reduced to a simple dynamic mean-field for a population averaged plastic synaptic current. Next, using the concepts of nonequilibrium thermodynamics, we derive the energy rate (entropy production rate) for plastic synapses and a corresponding Fisher information for coding presynaptic input. That energy, which is of chemical origin, is primarily used for battling fluctuations in the synaptic weights and presynaptic firing rates, and it increases steeply with synaptic weights, and more uniformly though nonlinearly with presynaptic firing. At the onset of synaptic bistability, Fisher information and memory lifetime both increase sharply, by a few orders of magnitude, but the plasticity energy rate changes only mildly. This implies that a huge gain in the precision of stored information does not have to cost large amounts of metabolic energy, which suggests that synaptic information is not directly limited by energy consumption. Interestingly, for very weak synaptic noise, such a limit on synaptic coding accuracy is imposed instead by a derivative of the plasticity energy rate with respect to the mean presynaptic firing, and this relationship has a general character that is independent of the plasticity type. An estimate for primate neocortex reveals that a relative metabolic cost of BCM type synaptic plasticity, as a fraction of neuronal cost related to fast synaptic transmission and spiking, can vary from negligible to substantial, depending on the synaptic noise level and presynaptic firing.

## 1. Introduction

Information and energy are intimately related for all physical systems because information has to be written on some physical substrate which always comes at some energy cost (Landauer 1961; Bennett 1982; Leff and Rex 1990; Berut et al 2012; Parrondo et al 2015). Brains are physical devices that process information and simultaneously dissipate energy (Levy and Baxter 1996; Laughlin et al 1998) in the form of heat (Karbowski 2009). This energetic cost is relatively high (Laughlin et al 1998; Aiello and Wheeler 1995; Attwell and Laughlin 2001; Karbowski 2007), which is the likely cause for a sparse coding strategy in neural circuits (Balasubramanian et al 2001; Niven and Laughlin 2008). Experimental studies (Shulman et al 2004; Logothetis 2008; Alle et al 2009), as well as theoretical calculations based on data (Harris et al 2012; Karbowski 2012), indicate that fast synaptic signaling, i.e. synaptic transmission, together with neuron’s action potentials are the major consumers of metabolic energy. This type of energy use is of electric origin, and is caused by flows of electric charge due to voltage and concentration gradients and subsequent pumping ions out to maintain these gradients (Attwell and Laughlin 2001; Karbowski 2009). This very pumping of electric charge requires large amounts of energy.

Brains are also highly adapting objects, which learn and remember by encoding and storing long-term information in excitatory synapses (dendritic spines) (Kasai et al 2003; Takeuchi et al 2014). These important slow processes are driven by correlated electric activities of pre- and post-synaptic neurons (Markram et al 1997; Bienenstock et al 1982; Miller and MacKay 1994, Song et al 2000; Van Rossum et al 2000) and cause plastic modifications in spine’s intrinsic molecular machinery, leading to changes in spine size, its conductance (weight) and postsynaptic density (PSD) (Kasai et al 2003; Bonhoeffer and Yuste 2002; Holtmaat et al 2005; Meyer et al 2014). Consequently synaptic plasticity and associated information writing and storing must cost energy, since spines require some energy for inserting and maintaining AMPA and NMDA receptors on spine membrane (Huganir and Nicoll 2013; Choquet and Triller 2013), as well as for powering various molecular processes associated with PSD (Lisman et al 2013; Miller et al 2005). In contrast to fast synaptic transmission and neuron discharges, which are of electric nature, the plastic slow synaptic processes are of chemical origin (interactions between spine proteins), and thus require the chemical energy (Karbowski 2019).

One of the empirical manifestations of the plasticity-energy relationship is present for mammalian cortical development, during which synaptic density can change several fold and strongly correlates with changes in glucose metabolic rate of cortical tissue (Karbowski 2012). Unfortunately, despite a massive literature on modeling synaptic plasticity (e.g. Bienenstock et al 1982; Miller and MacKay 1994, Song et al 2000; Van Rossum et al 2000; Billings and Van Rossum 2009; Clopath et al 2010; Pfister and Gerstner 2006; Tetzlaff et al 2011; Toyoizumi et al 2014; Costa et al 2015; Shouval et al 2002; Graupner and Brunel 2012; Ziegler et al 2015; Fusi et al 2005; Benna and Fusi 2016; Gutig et al 2003; Smolen et al 2012), our theoretical understanding of the energetic requirements underlying synaptic plasticity and memory storing is currently lacking. In particular, we do not know the answers to the basic questions, such as how does energy consumed by plastic synapses depend on key neurophysiological parameters, and more importantly, whether energy restricts the precision of synaptically encoded information and its lifetime, and to what extent. Such a knowledge might lead to a deeper understanding of two fundamental problems in neuroscience: one related to the physical cost and control of learning and memory in the brain (Kasai et al 2003; Takeuchi et al 2014; Lisman et al 2013; Costa et al 2015; Kandel et al 2014; Chaudhuri and Fiete 2016; Zenke and Gerstner 2017), and another more practical related to dissecting the contribution of synaptic plasticity to signals in brain imaging (Attwell and Laughlin 2001; Logothetis 2008; Engl et al 2017; Magistretti et al 1999; Shulman et al 2004). A recent study by the author (Karbowski 2019) provided some answers to the above questions, by analyzing molecular data in synaptic spines and by modeling energy cost of learning and memory in a cascade model of synaptic plasticity (mimicking molecular interactions in spines). From that study it follows that the average cost of synaptic plasticity constitutes a small fraction of the metabolic cost used for fast excitatory synaptic transmission, about 4 − 11%, and that storing longer memory traces can be relatively cheap (Karbowski 2019). However, this study left open other questions, e.g., how does the energy cost of synaptic plasticity depend on neuronal firing rates, synaptic noise, and other neural characteristics, and what is the relationship between such energy cost and a precise storing of synaptic information?

The main goal of this study is to uncover a relationship between synaptic plasticity, its energetics, and a precise information storing at excitatory synapses for one of the best known forms of synaptic plasticity due to Bienenstock, Cooper, and Munro, the so-called BCM rule (Bienenstock et al 1982). This is a different (more macroscopic) but a complementary level of modeling to the one (microscopic) in (Karbowski 2019). Specifically, we want to determine the energy cost associated with the accuracy of information encoded in a population of plastic synapses about the presynaptic input. Additionally, we want to find the relationship between plastic energy rate and memory duration about a single event at synapses. In other words, our goal is to find the metabolic requirement of maintaining an accurate information at synapses in the face of ongoing variable neural activity and thermodynamic fluctuations inside spines associated with variation in the number of membrane receptors. The phenomenological BCM rule has been shown to explain several key experimental observations (Cooper and Bear 2012), and it is equivalent to a more microscopic STDP rule (Markram et al 1997; Song et al 2000; Van Rossum 2000) under some very general conditions (Pfister and Gerstner 2006; Izhikevich and Desai 2003). Since, the BCM rule is believed to describe initial phases of learning and memory (Zenke and Gerstner 2017), the focus of this work is on the energy cost and coding accuracy of the early synaptic plasticity, i.e. early long-term potentiation (e-LTP) and depression (e-LTD), which lasts from minutes to several hours. We do not consider explicitly the effects of memory consolidation that operate on much longer time scales and which are associated with late phases of LTP and LTD (l-LTP and l-LTD) (Ziegler et al 2015; Benna and Fusi 2016; Redondo and Morris 2011). However, we do provide a rough estimate of the energetics of these late processes, and they turn out to be much less energy demanding than the early phase plasticity.One can question whether the approach taken here, with the macroscopic BCM type model, is reasonable for modeling and calculating energy cost of synaptic plasticity? Maybe a more microscopic approach should be used with explicit molecular interactions between PSD proteins? However, the basic problem with such a microscopic more detailed approach is that we do not know most of the molecular signaling pathways in a dendritic spine, we do not know the rates of various reactions, and even the basic mechanism of encoding information at synapses is unclear. For example, for a long time it was thought that CaMKII persistent autophosphorylation provides a basic mechanism of information storage via bistability (Lisman et al 2013; Miller et al 2005). However, experimental data indicate that CaMKII enhanced activity after spine activation is transient and lasts only about 2 min (Lee et al 2009), which casts doubts on its persistent enzymatic activity and its role as a “memory molecule” (for a review see, Smolen et al 2019). Taking all these uncertainties into account, it seems that more macroscopic approach might be more reliable, at least partly.

Because synapses/spines are small, they are strongly influenced by thermal fluctuations (Kasai et al 2003; Choquet and Triller 2013; Statman et al 2014). For this reason, this paper uses universal methods of stochastic dynamical systems and non-equilibrium statistical mechanics (Nicolis and Prigogine 1977; Van Kampen 2007; Risken 1996; Lan et al 2012; Mehta and Schwab 2012; Tome 2006; Tome and de Oliveira 2010; Seifert 2012). The latter are generally valid for all physical systems, including the brain, operating out of thermodynamic equilibrium. Regrettably, the methods of non-equilibrium thermodynamics have virtually not been used in neuroscience despite their large po-tential in linking brain physicality with its information processing capacity, with two recent exceptions (Goldt and Seifert 2017; Karbowski 2019). (This should not be confused with equilibrium thermodynamics, whose methods have occasionally been used in neuroscience, although in a different context, e.g., Balasubramanian et al 2001; Tkacik et al 2015; Friston 2010.)

### General outline of the problem considered

It is generally believed that long-term information in excitatory synapses is encoded in the pattern of synaptic strengths or weights (membrane electric conductance), which is coupled to the molecular structure of postsynaptic density within dendritic spines (Takeuchi et al 2014; Lisman et al 2013; Miller et al 2005; Kandel et al 2014, Zhu et al 2016). This study considers the energy cost associated with maintaining the pattern of synaptic weights. In particular, we analyze the energetics and information capacity of the fluctuations in the number of AMPA and NMDA receptors on a spine membrane, or equivalently, fluctuations in the synaptic conductance. Such a variability in the receptor number tends to spread the range of synaptic weights (affecting their structure and distribution) that has a negative consequence on the encoded information and can lead to its erasure. In terms of statistical mechanics, the receptor fluctuations increase the entropy associated with the distribution of synaptic weights, and that entropy has to be reduced to preserve the information encoded in the weights. This very process of reducing the synaptic entropy production is a nonequilibrium phenomenon that costs some energy, which has to be provided by various processes involving ATP generation (Nicolis and Prigogine 1977).The BCM type of synaptic plasticity used here is a phenomenological model that does not relate in a straightforward way to the underlying synaptic molecular processes. Empirically speaking, a change in synaptic weight in e-LTP is caused by a sequence of molecular events, of which the main are: activation of proteins in postsynaptic density, which subsequently stimulates downstream actin filaments elongation (responsible for a spine enlargement), and AMPA and NMDA receptor trafficking (Huganir and Nicoll 2013; Choquet and Triller 2013). Therefore, it is assumed here that BCM-type rule used here macroscopically reflects broadly these three microscopic processes, especially the first and the last. (Spine volume related to actin dynamics is not explicitly included in the model, although it is known experimentally that spine volume and conductance are positively correlated (Kasai et al 2003).) Thus, it is expected that the synaptic energy rate calculated here is related to ATP used mainly for postsynaptic protein activation through phosphorylation process (Zhu et al 2016), and receptor insertion and movement along spine membrane. Obviously, there are many more molecular processes in a typical spine, but they are either not directly involved in spine conductance variability or they are much faster than the above processes (e.g. releasing Ca^2+^ from internal stores is fast). A detailed empirical estimation based on molecular data suggests that protein activation via enhanced phosphorylation is the dominant contribution to the energy cost (ATP rate) of synaptic plasticity (Karbowski 2019). Therefore, the theoretical energy rate of synaptic plasticity determined here should be viewed as a minimal but a reasonable estimate of energetic requirement of LTP and LTD, and it is strictly associated with the information encoded in synaptic weights.Experimental data show that excitatory synapses can exist in two or more stable states, characterized by discrete synaptic weights or sizes (Kasai et al 2003; Montgomery and Madison 2004; Petersen et al 1998; O’Connor et al 2005; Loewenstein et al 2011; Bartol et al 2015). Data on a single synapse level indicate that synapses can operate as binary elements either with low or high electric conductance (Petersen et al 1998; O’Connor et al 2005). On the other hand, the data on a population level, more relevant to this work, show that synapses can assume more than two stable discrete states (Kasai et al 2003; Loewenstein et al 2011; Bartol et al 2015). In either case, the issue of bistability vs. multistability is not yet resolved. In this study, a minimal scenario is considered in which synapses together with their postsynaptic neuron can effectively act as a binary coupled system, characterized by a single variable, which is the meanfield postsynaptic current with one or two stable states. The bistability is produced here from an extended BCM model, which in principle allows for continuous changes in synaptic weights for individual synapses. The important point is that these continuous weights are correlated, due to plasticity constraints, and thus converge on a mean-field population level either to one or to a couple of stable values.

Synaptic plasticity processes are induced by a correlated firing in pre- and post-synaptic neurons, and thus a model of neuron activity is also needed. This study uses a firing rate neuron model of the so-called class one nonlinear firing rate curve, which is believed to be a good approximation to biophysical neuronal models (Ermentrout 1998; Ermentrout and Terman 2010); see the Methods for details.

The paper is organized as follows. First, we introduce and solve an extended model of the classical BCM plasticity rule. Then, we derive an effective equation for the meanfield stochastic dynamics of the synaptic currents for that extended plasticity model. Next, we translate this effective equation into probabilistic Fokker-Planck formalism, and derive an effective potential for the mean-field synaptic current. With the help of the effective potential we find entropy production and Fisher information associated with the synaptic plasticity stochastic dynamics. Entropy production is related to the energy cost of the extended BCM plasticity, while the Fisher information is related to the accuracy of encoded information in a population of plastic synapses about the presynaptic input. Details of the calculations are provided in the Methods (and some in Supporting Information S1).

## 2. Results

### Model of synaptic plasticity: stochastic BCM type

We consider a sensory neuron with *N* plastic excitatory synapses (dendritic spines). We assume that synaptic weights *w_i_* (*i* = 1,…, *N*), corresponding to spine electric conductances, change due to two factors: correlated activity in presynaptic and postsynaptic firing rates (*f_i_* and *r*, respectively), and noise in spine conductance (~ *σ_w_*). The noise is caused by two basic factors: an internal thermodynamic fluctuations in spines because of their small size (< 1 μm) and relatively small number of molecular components (Kasai et al 2003; Statman et al 2014), and by presynaptic fluctuations in the firing rates that drive the ionic and molecular fluxes in spines. The dynamics of synaptic weights is given by a modified BCM plasticity rule (Bienenstock et al 1982):

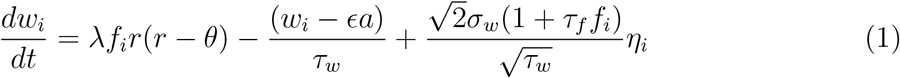

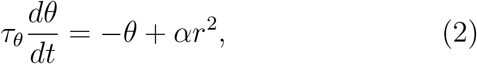

where λ is the amplitude of synaptic plasticity controlling the rate of change of synaptic conductance, *τ_w_* is the weights time constant controlling their decay duration, *θ* is the homeostatic variable the so-called sliding threshold (adaptation for plasticity) related to an interplay of LTP and LTD with time constant *τ_θ_*, and *α* is the coupling intensity of *θ* to the postsynaptic firing rate *r*. The noise term in Eq. (1) is represented as Gaussian white noise *η_i_* with zero mean and Delta function correlations, i.e., 〈*η_i_*(*t*)〉_*η*_ = 0 and 〈*η_i_*(*t*)*η_j_*(*t*’))_*η*_ = *δ_ij_δ*(*t* – *t*’) (van Kampen 2007). The amplitude of the noise in weights is proportional to the standard deviation *σ_w_* (in units of conductance) due to basic thermodynamic fluctuations in spines, and to the factor (1 + *τ_f_f_i_*) due to additional fluctuations in the presynaptic activities. The latter factor simply amplifies the basic thermodynamic fluctuations. The time scale *τ_f_* of fluctuations in *f_i_* was added in the noise term to maintain a unitless form of the amplifying factor. Finally, the product *ϵa* is the minimal synaptic weight when there is no presynaptic stimulation (*f_i_* = 0), where the unitless parameter *ϵ* ≪ 1. There are two modifications to the conventional BCM rule: the stochastic term ~ *σ_w_*, and the decay term of synaptic weights with the time constant *τ_w_*, which is key for reproducing a binary nature of synapses (Petersen et al 1998; O’Connor et al 2005) and for determining energy used by synaptic plasticity.

The conventional BCM rule (i.e. for *τ_w_* Ⅺ ∞ and *σ_w_* = 0) describes temporal changes in synaptic weights due to correlated activity of pre- and post-synaptic neurons (both *f_i_* and *r* are present on the right in Eq. (1)). These activity changes can either increase the weight, if postsynaptic firing *r* is greater than the sliding threshold *θ* (this corresponds to LTP), or they can decrease the weight if *r* < *θ* (corresponding to LTD). The interesting aspect is that *θ* is also time dependent, and it responds quickly to changes in the postsynaptic firing. In effect, when both dynamical processes in Eqs. (1–2) are taken into account, the synapse is potentiated for low *r* (LTP) and depressed for high *r* (LTD).

We assume, in accordance with empirical data, that presynaptic firing rates *f_i_* change on a much faster time scale (*τ_f_* ~ 0.1 − 1 sec) than the synaptic weights *w_i_* (changes on time scale *τ_w_* ~ 1 hr). We further assume that each presynaptic firing rate *f_i_* fluctuates stochastically around mean value *f_o_* with a standard deviation *σ_f_*, and that these fluctuations are uncorrelated. This implies that there is a time scale separation between neural activities and synaptic plasticity activities.

### Numerical solution of the stochastic extended BCM plasticity model

In this section we solve numerically the model represented by *N* +1 Eqs. (1–2).

We first consider the model without synaptic noise, i.e., *σ_w_* = 0. This deterministic system can exhibit collective bistability, regardless of whether *σ_f_* is 0 or finite. The critical factor in generating bistability is that the time constant *τ_θ_* for the homeostatic variable *θ* is much smaller than synaptic plasticity time constant *τ_w_*. That is, the variable *θ* must be much faster than the synaptic weights *W_i_*. Typically, bistability is found for *τ_θ_/τ_w_* ≤ 0.06, and we work in this regime throughout the whole study (for the neurobiological validity of this regime, see the Discussion section). Collective bistability means that all synaptic weights can converge to two different fixed points depending on the initial conditions (Fig. 1). When all synapses start from sufficiently small weights, then they all converge into the same small synaptic weight *ea.* If the initial weights are much larger than ea, then all synapses become asymptotically strong. Thus, there is a strong collective behavior of synapses in the deterministic case.

**Fig. 1.**
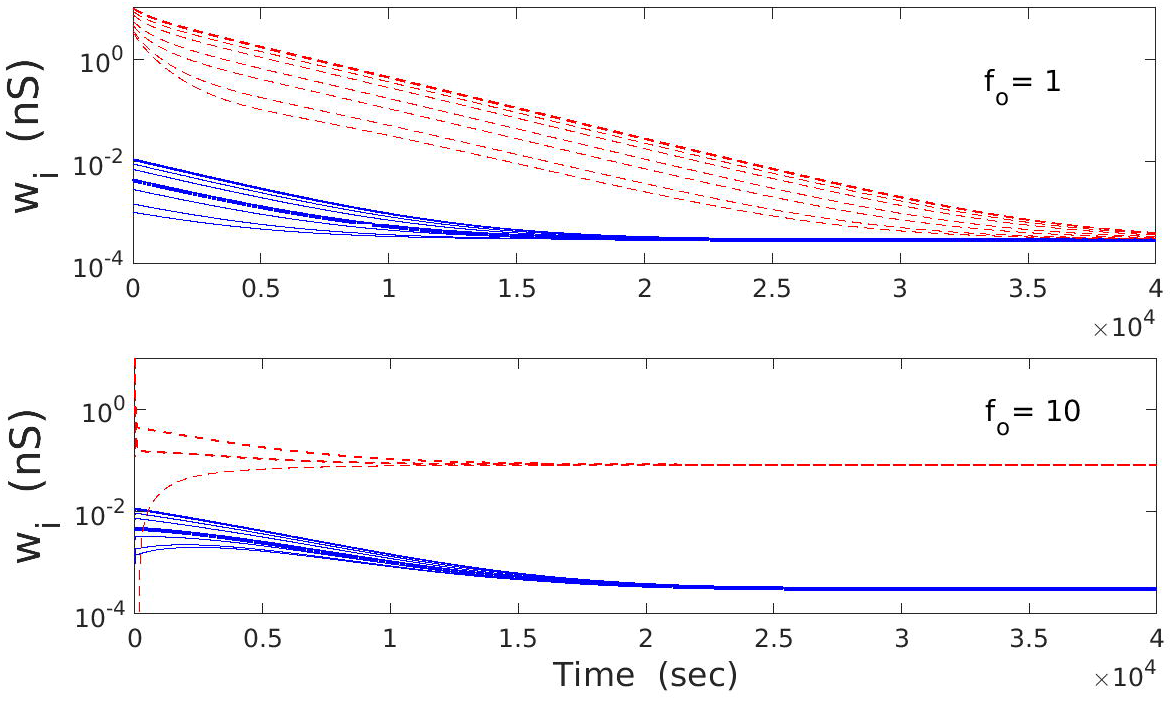
Asymptotic behavior of the deterministic extended BCM model. Temporal dependence of deterministic synaptic weights *w_i_* from Eqs. (1–2). (Upper panel) When mean firing rate *f_o_* is smaller than some threshold value, then all weights converge on the same asymptotic value regardless of the initial values (red lines for strong initial synapses, and blue lines for weak initial synapses). This case corresponds to monostability. (Lower panel) When *f_o_* is in the intermediate interval, then synaptic weights can converge into two separate values, depending on initial conditions. In this regime, there are two different coexisting fixed points (bistability). Strong initial weights lead to large final value (red lines), and weak initial weights lead to low final value (blue lines). Parameters used: λ = 9 · 10^−7^ and *κ* = 0.001.

Two main parameters that control the shape of the bifurcation diagram are the synaptic plasticity amplitude λ and the mean firing rate *f_o_* (Fig. 2). For a fixed *f_o_*, synapses can be either in monostable or bistable phase, depending on the value of λ (Fig. 2A). Generally, for small and sufficiently large λ there is monostability, while for intermediate λ there is bistability. For a fixed λ, the picture is slightly more complex: bistability can emerge already for *f_o_* = 0 (for intermediate λ), or for some finite *f_o_* (for small *λ*), or bistability can never appear (for large *λ*) (Fig. 2A). The bifurcation diagram, i.e., the dependence of asymptotic value of *w_i_* on *f_o_* is presented in Fig. 2B.

**Fig. 2.**
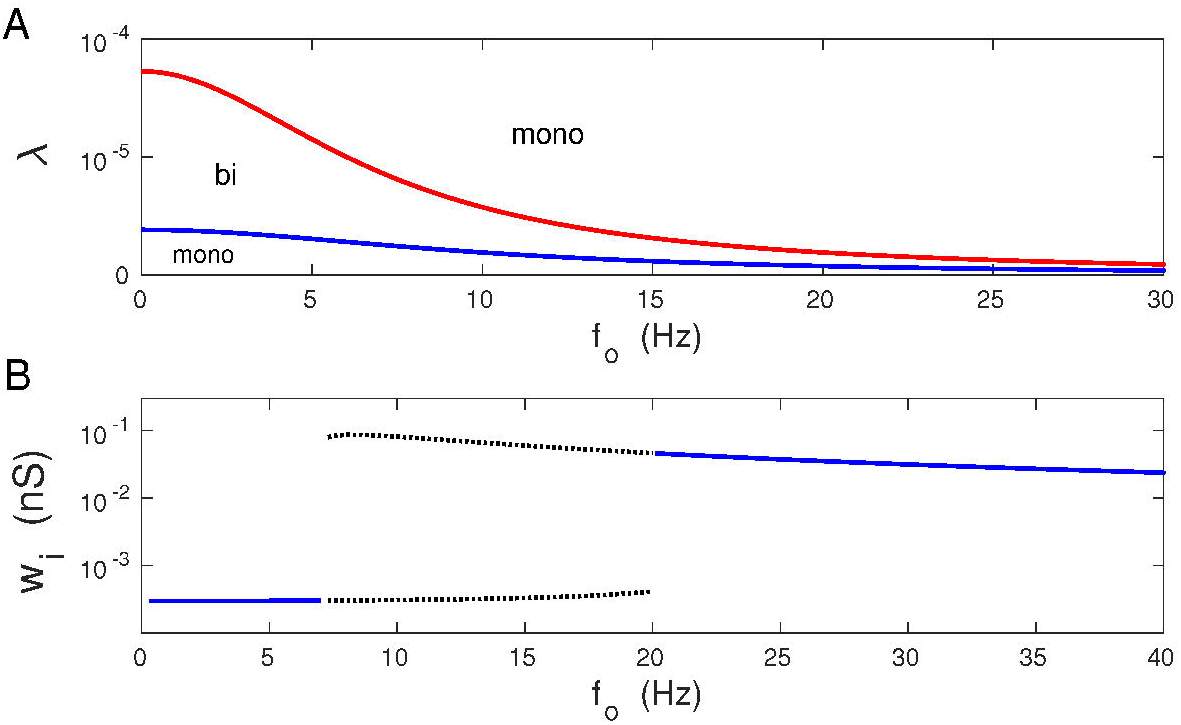
Phase and bifurcation diagrams of *w_i_* for the deterministic extended BCM model. (A) Schematic phase diagram λ vs. *f_o_* for monostable (mono) and bistable (bi) behavior of synaptic weights *w_i_*. Bistability is lost for sufficiently large λ. Note that the critical values of λ and *f_o_*, for which bistability emerges and disappears, are inversely related. (B) Bifurcation diagram of an asymptotic synaptic weight vs. *f_o_* for a typical synapse. The bistable regime is indicated by dotted lines. Parameters used: λ = 9 · 10^−7^, and *κ* = 0.001.

Inclusion of synaptic noise, i.e. *σ_w_* > 0, leads to stochastic fluctuations of individual synapses. In the monostable regime, fluctuations are around a given fixed point (either weak or strong weight). In the bistable regime, individual synapses fluctuate between weak and strong weights (Fig. 3A). Despite synaptic noise, the collective nature of synapses is statistically preserved, as all synapses have similar weight distributions (Fig. 3B, C). These distributions are much more spread in the bistable regime than in the monostable, and they seem to be almost uniform for bistability.

**Fig. 3.**
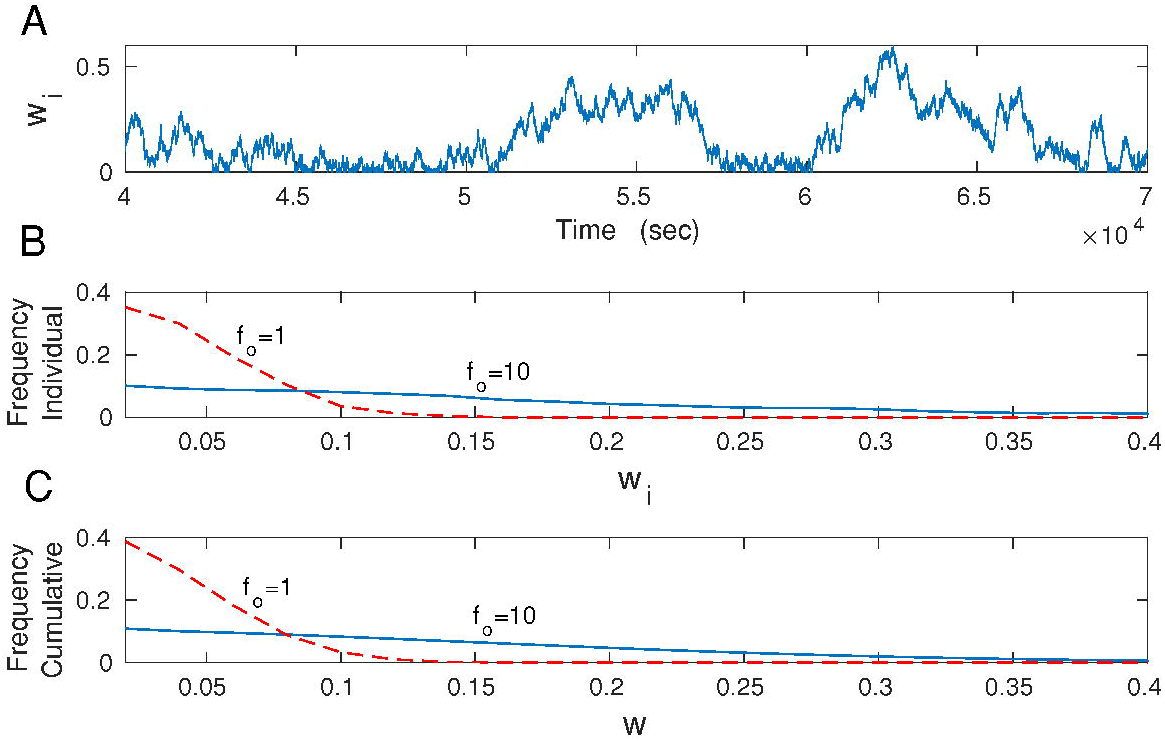
Distributions of synaptic weights *w_i_* for the stochastic extended BCM model. (A) Temporal fluctuations of an individual synapse in the bistable regime. Note stochastic jumps between weak and strong weights. (B) Distribution of synaptic weights for an individual synapse (the same as in A) in the bi-(*f_o_* = 10 Hz; blue solid line) and monostable (*f_o_* = 1 Hz; red dashed line) regimes. (C) Cumulative distribution of synaptic weights for all *N* synapses in the bi-(*f_o_* = 10 Hz; blue solid line) and mono-stable (*f_o_* = 1 Hz; red dashed line) regimes. Note that both distributions in (B) and (C) are very similar. Parameters used: λ = 9 · 10^−7^, *κ* = 0.001, and *σ_w_* = 0.1 nS.

### Dimensional reduction of the stochastic BCM model: dynamic mean field

The stochastic system of *N* +1 equations described by Eqs. (1–2) is not tractable analytically, because it is the coupled nonlinear system. The coupling takes place via postsynaptic firing rate r, which depends on all synaptic weights *w_i_* (in Eqs. 1–2). In this section an effective mean-field model corresponding to the extended BCM model in Eqs. (1–2) is presented and discussed that is amenable to analytical considerations. In this dynamical mean-field, we focus on a single dynamic variable, which is a population averaged synaptic current *v* defined by Eq. (25) in the Methods. The single variable *v* is sufficient to describe the global stochastic dynamics of the original model given by Eqs. (1–2), because together with postsynaptic firing *r* it forms a closed mathematical system of just two equations; see below. The practical reason behind introducing the dynamic mean-field is that this approach enables us to obtain explicit formulae for synaptic plasticity energy rate and coding accuracy.

We can reduce the multidimensional system (1-2) into a single effective equation, primary because of the time scale separation between neural firing dynamics (changes typically on the order of seconds or less) and between synaptic plasticity (changes on the timescale of minutes/hours). Moreover, we assume that the two synaptic plasticity processes, described by Eqs. 1 and 2, have two distinct time scales, and *τ_w_* dominates over *τ_θ_* in duration. For the neurophysiological validity of this assumption, see the Discussion. Consequently, for times of the order of *τ_w_*, we have *dθ/dt* ≈ 0, which implies that *θ* ≈ *αr*^2^. The details of the reduction procedure can be found in the Methods, in which we obtain a single plasticity equation for a population averaged excitatory postsynaptic current *v* per synapse, which is related to *w_i_* and *f_i_* by *v* = (*β/N*) ∑_*i*_ *f_i_w_i_*, where *β* depends on neurophysiological parameters and is defined in Eq. (25). The result of the reduction procedure is

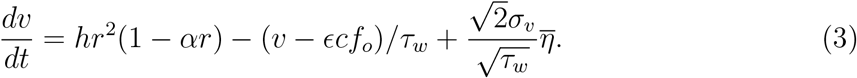

This equation essentially couples slow synaptic activities with fast neural activities, and gives a single equation describing the mean-field dynamics of the coupled system: synapses plus their postsynaptic neuron. In Eq. (3), the symbol *h* is the drivingplasticity parameter given by

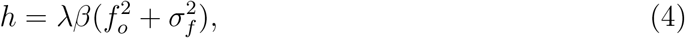

with *f_o_* and *σ_f_* denoting the mean and standard deviation of presynaptic firing rates. Mathematically, the driving-plasticity *h* is proportional to the product of plasticity amplitude λ and the presynaptic driving 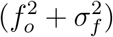, which implies that *h* grows quickly with the presynaptic firing rate. Physically, *h* is proportional to the electric charge that, on average, can enter the spine due to a correlated activity of pre- and post-synaptic neurons (*h* has a unit of electric charge). This means that the magnitude of *h* is a major determinant of the plasticity (driving force counteracting the synaptic decay), since larger *h* can experimentally correspond to more Ca^+2^ entering the spine and a higher chance of invoking a change in synaptic strength, which agrees qualitatively with the experimental data (Huganir and Nicoll 2013; Lisman et al 2012).

The rest of the parameters in Eq. (3) are *c* = *aβ*, and 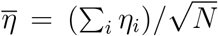, which denotes a new (population averaged) Gaussian noise with zero mean and delta function correlations. This population noise has the amplitude *σ_v_*, which corresponds to a standard deviation of *v* when *h* = 0, and it is given by

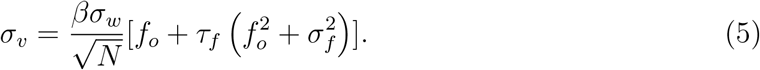

Note that *σ_v_* scales as 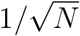, and it is a product of the intrinsic synaptic conductance noise and of the presynaptic neural activity. The latter implies that a higher presynaptic activity amplifies the current noise.

In Eq. (3), the postsynaptic firing rate *r* assumes its quasi-stationary value (due to time scale separation), and is related to v through (for details see the Methods):

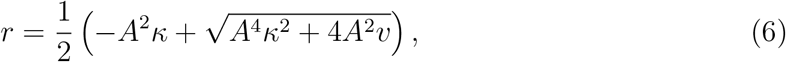

where *A* is the postsynaptic firing rate amplitude, and *κ* is the intensity of firing rate *r* adaptation. Broadly speaking, the magnitude of *κ* reflects the strength of neuronal self-inhibition due to adaptation to synaptic stimulation (see Eqs. 21 and 22 in the Methods). Generally, increasing *κ* leads to decreasing postsynaptic firing rate *r* (Fig. 4A). For *κ* = 0, we recover a nonlinear firing rate curve (square root dependence on synaptic current *v*) that is characteristic for class one neurons (Ermentrout 1998; Ermentrout and Terman 2010), while for sufficiently large *κ*, i.e. for 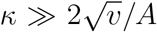, we obtain a linear firing rate curve *r*(*v*) ≈ *v/κ* (Fig. 4A). Equations (3) and (6) form a closed system for determining the stochastic dynamics of the postsynaptic current *v*.

**Fig. 4.**
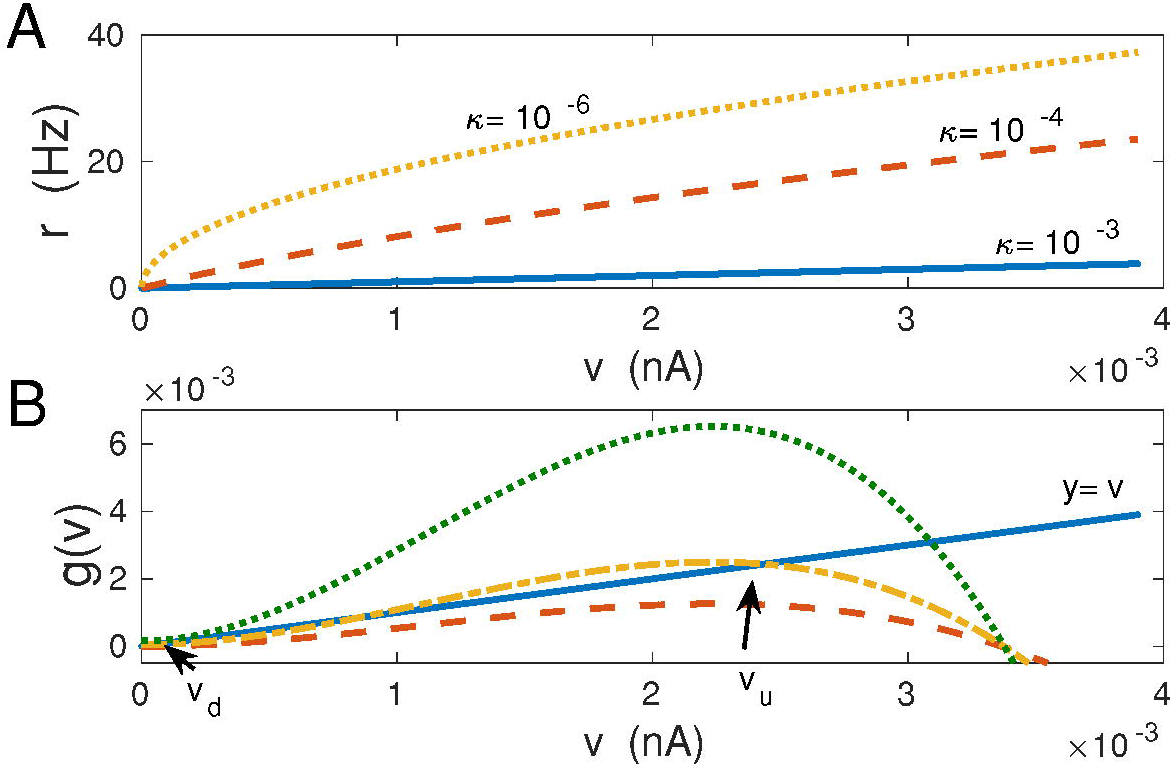
Firing rates and emergence of bistability in the mean-field model: theory. (A) Postsynaptic firing rates *r* as functions of the population averaged synaptic current *v* for different neuronal adaptations values *κ* (in nA·sec). Increasing *κ* causes decrease in *r* and makes the functional form *r*(*v*) more linear. (B) Graphical solutions of Eq. (7) and multiple roots for stationary *v*. For small driving-plasticity *h* there is only one intersection of *g*(*v*) and the line *y* = *v* at *v* ~ *O*(*ϵ*), corresponding to *v_d_* and monostability (dashed red line; *f*_0_ = 0.3 Hz, *σ_f_* = 8 Hz). For higher *h* (*h* > *h_cr_*) there are three intersections, but the middle one corresponds to an unstable solution, which in effect yields two stable solutions, i.e. bistability (dashed-dotted yellow line; *f*_0_ = 5.0 Hz, *σ_f_* = 10 Hz). When *h* is very large, then there is only one intersection, and it occurs for large *v*, corresponding to monostability with strong synapses *v_u_* only (dotted green line; *f*_0_ = 30.0 Hz, *σ_f_* = 10 Hz). In panel (B) λ = 9 · 10^−7^ and *κ* = 0.001.

### Geometric steady state solution of the deterministic mean-field: emergence of bistability in *v*

We can use geometric considerations to gain some intuitive understanding of the meanfield deterministic behavior represented by Eqs. (3) and (6). If we put *dv/dt* = 0 and *σ_v_* = 0 in Eq. (3), we can rearrange it to obtain

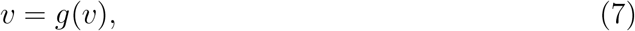

where the right hand side of this equation, *g*(*v*) = *ϵcf_o_* + *τ_w_hr*^2^(1 − *αr*), and it depends on *v* only through *r* as in Eq. (6) (see Fig. 4A). Moreover, the function *g*(*v*) has a maximum with height proportional to *h*. When *h* is very small, Eq. (7) has only one solution *v* ~ *O*(*e*) (i.e. one intersection point of the curves representing the functions on the right and on the left; Fig. 4B). This solution corresponds to weak synapses and monostable regime. Increasing *h*, by increasing *f*_0_, causes an increase in the maximal value of the right hand side in Eq. (7), such that more solutions are possible (Fig. 4B). In particular, when *h* grows above a certain critical value *h_cr_*, Eq. (7) generates 3 solutions (one ~ *O*(*ϵ*) and two other ~ *O*(1)), of which the middle one is unstable (Fig. 4B). This case corresponds to bistable regime with two stable solutions, representing weak and strong synaptic currents that can be called, respectively, “down” and “up” synaptic states. These two states could hypothetically be related to thin and mushroom dendritic spines, with small and large number of AMPA receptors, respectively (Bourne and Harris 2007). For very large driving-plasticity *h* the two lower solutions disappear and we have again a monostable regime with strong synapses only (Fig. 4B).

A geometrical condition for the emergence of bistability is when the function *g*(*v*) in Eq. (7) first touches tangentially the line *y* = v, i.e. when *dg/dv* = 1 (Fig. 4B). Solving this condition together with Eq. (6) yields for *ϵ* ≪ 1 the critical value of the driving-plasticity parameter hcr as

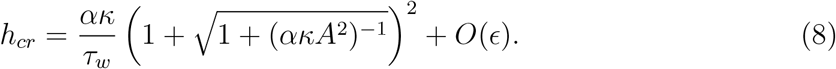

Note that for very fast decay in Eq. (3), i.e. for *τ_w_* ↦ 0, the bistability is lost, since then *h_cr_* ↦ ∞, and there is only one solution corresponding to weak synapses *v* ~ *O*(*ϵ*). Bistability is also lost in the opposite limit of extremely slow decay, *τ_w_* ↦ ∞, but in this case the only stable solution corresponds to strong synapses. Interestingly, for very strong neural adaptation, *κ* ↦ ∞, the bistability also disappears, since then *h_cr_* ⊦ ∞. This case corresponds to extremely small postsynaptic firing rates, *r* ≈ *v/κ* ≈ 0 (Fig. 4A), and indicates the absence of a driving force capable of pushing synapses to a higher conducting state. On the other hand, when there is no adaptation, *κ* → 0, the critical *h_cr_* ↦ (*τ_w_A*^2^)^−1^, i.e. it is finite. This means that it is easier to produce synaptic bistability for neurons with stronger nonlinearity in their firing rate curves (see Eq. 6; Fig. 4A).

### Analytic steady state solution of the deterministic mean-field

The above geometric intuition can be supported by an analytic approach. In the deterministic limit, *σ_v_* = 0 (which is obtained either for *N* → ∞ or for *σ_w_* = 0), we can solve the mean-field model of Eq. (3) in the stationary state, i.e. we can find its fixed points by setting *dv/dt* = 0. To achieve this, it is more convenient to work with the postsynaptic firing rate *r* than with *v* variable, due to a nonlinear dependence of *r* on *v*, which is given by Eq. (6). Inverting Eq. (6), we find *v* = *κr* + (*r/A*)^2^, which can be used in the condition *dv/dt* = 0. This generates an equation for roots of the cubic polynomial in the *r* variable:

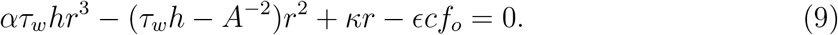

The discriminant Δ of this equation is

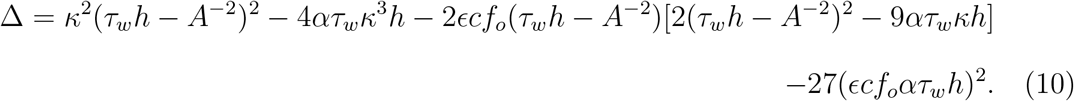

The sign of Δ determines how many real roots Eq. (9) has. Specifically, if Δ < 0, then Eq. (9) has one real root, whereas if Δ > 0 then Eq. (9) has three distinct real roots. The former case corresponds to monostability, while the latter to bistability in the mean-field deterministic dynamics of Eq. (3). The transition between this two regimes takes place for Δ = 0. The existence of these regimes obviously depends on the values of various parameters in the discriminant Δ.

The phase diagram of mono-vs. bi-stability in the parameter space of λ, *f_o_* (plasticity amplitude vs. mean presynaptic firing) is shown in Fig. 5. A specific bifurcation diagram, computed numerically from Eq. (3) for *σ_v_* = 0, in which a stationary value of *v* is plotted as a function of *f_o_* is presented in Fig. 6. The phase and bifurcation diagrams for stationary *v* in Figs. 5 and 6 look qualitatively similar to the phase and bifurcation diagrams of stationary *w_i_* in Fig. 2 for the full *N*-synaptic system of Eqs (1-2).

**Fig. 5.**
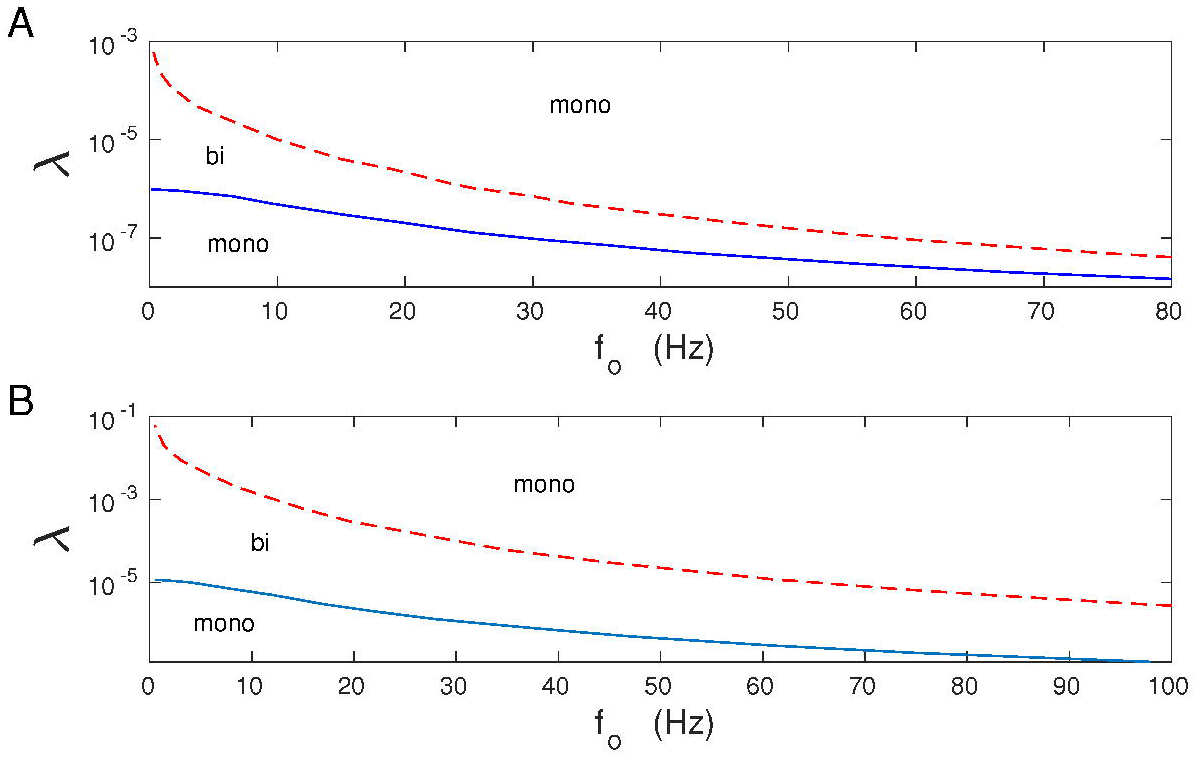
Phase diagram of mono-vs. bi-stability in the deterministic mean-field model. Numerical solution of Eq. (3) and (6) for *σ_v_* = 0. (A) *κ* = 0.001, (B) *κ* = 0.012. The solid and dashed lines are the boundaries between mono-(mono) and bistable (bi) regimes, and they correspond to the condition Δ = 0 in the (λ, *f_o_*) space.

**Fig. 6.**
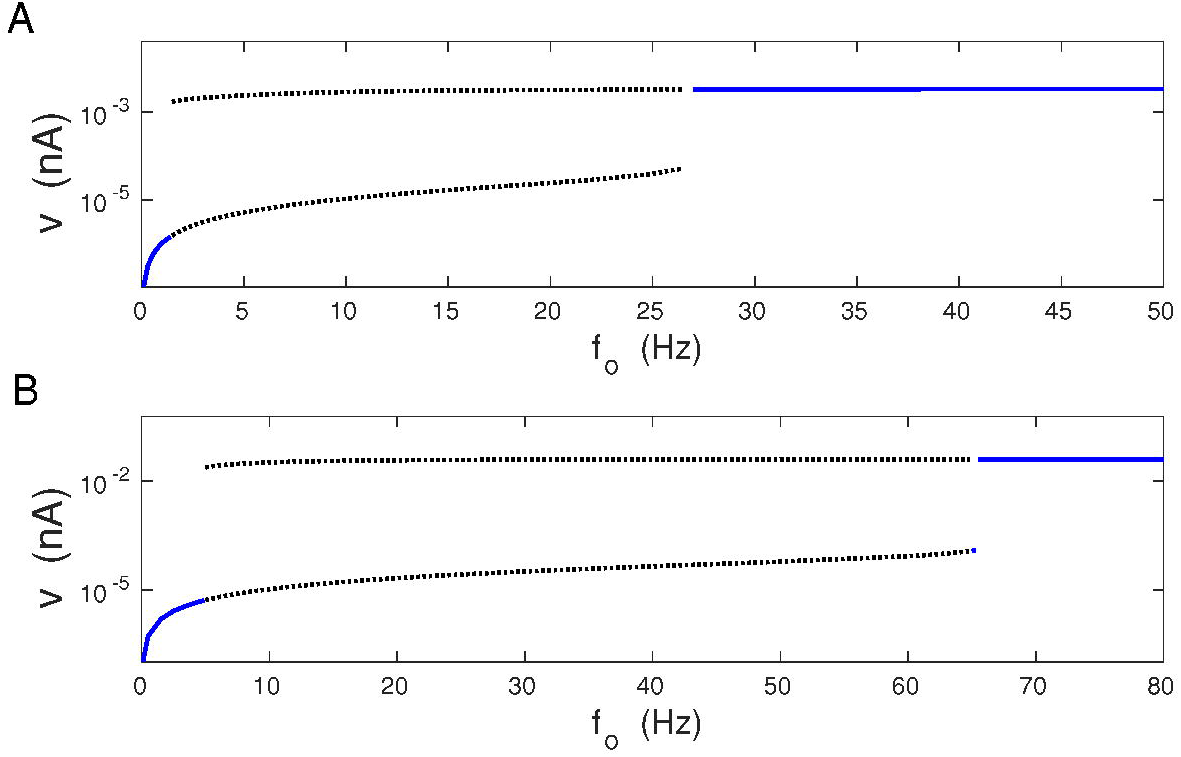
Bifurcation diagram of *v* in the deterministic mean-field model. Numerical solution of Eq. (3) and (6) for *σ_v_* = 0. (A) Parameters used: λ = 9 · 10^−7^ and *κ* = 0.001. (B) Parameters used: λ = 1 · 10^−5^ and *κ* = 0.012. Note that both diagrams in A and B are qualitatively similar, and there exists a critical value of *f_o_* above which bistability emerges.

Equation (9) can be solved analytically for small *ϵ*, as a series expansion in *ϵ*. The detailed procedure is described in the Suppl. Information S1. Depending on the sign of 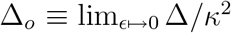, there can be one or three fixed points, which have the following values

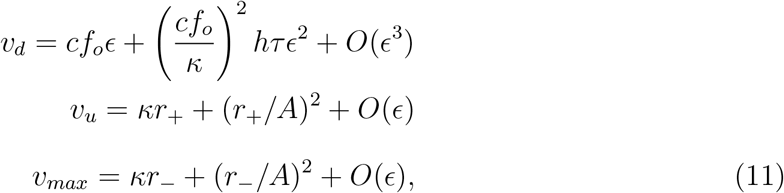

where

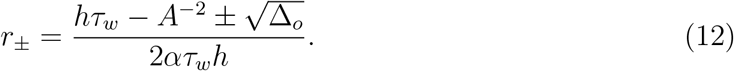

The value *v_d_* is the fixed point for weak synapses (down state), while *v_u_* is the fixed point for strong synapses (up state). The intermediate value *v_max_* corresponds to an always unstable fixed point, which serves as a boundary between the domains of attraction for down and up fixed points. Thus all initial values of v in the (0,*v_max_*) interval converge asymptotically into *v_d_*, and all initial values of *v* in the (*v_max_*, ∞) interval converge asymptotically into *v_u_*. From Eqs. (11) and (12) it can be seen that the value of the intermediate point *v_max_* decreases as *f_o_* (or *h*) increases, from the value *v_u_* (at the onset of bistability for Δ_*o*_ = 0) to the value *v_d_*. This mean that the domain of attraction for the *v_u_* fixed point increases at the expense of the domain of attraction for the *v_d_* fixed point, which shrinks with increasing *f_o_* in the bistable regime.

The critical value of the driving-plasticity parameter *h_cr_* for the emergence of bistability in Eq. (9) can be also obtained directly from the discriminant Δ in the limit *ϵ* ↦ 0. We obtain *h_cr_* by setting Δ_0_ = 0, and solving this equation for *h*.

### Stochastic mean-field: numerics and effective potential for synaptic current

When the synaptic noise is present, *σ_v_* > 0, the synaptic current v fluctuates. The distribution of *v* is unimodal for small mean firing rate *f_o_*, and bimodal for sufficiently large *f_o_* (Fig. 7). The bimodal distribution reflects the bistability found for the deterministic case, and it corresponds to synapses changing weights from weak to strong.

**Fig. 7.**
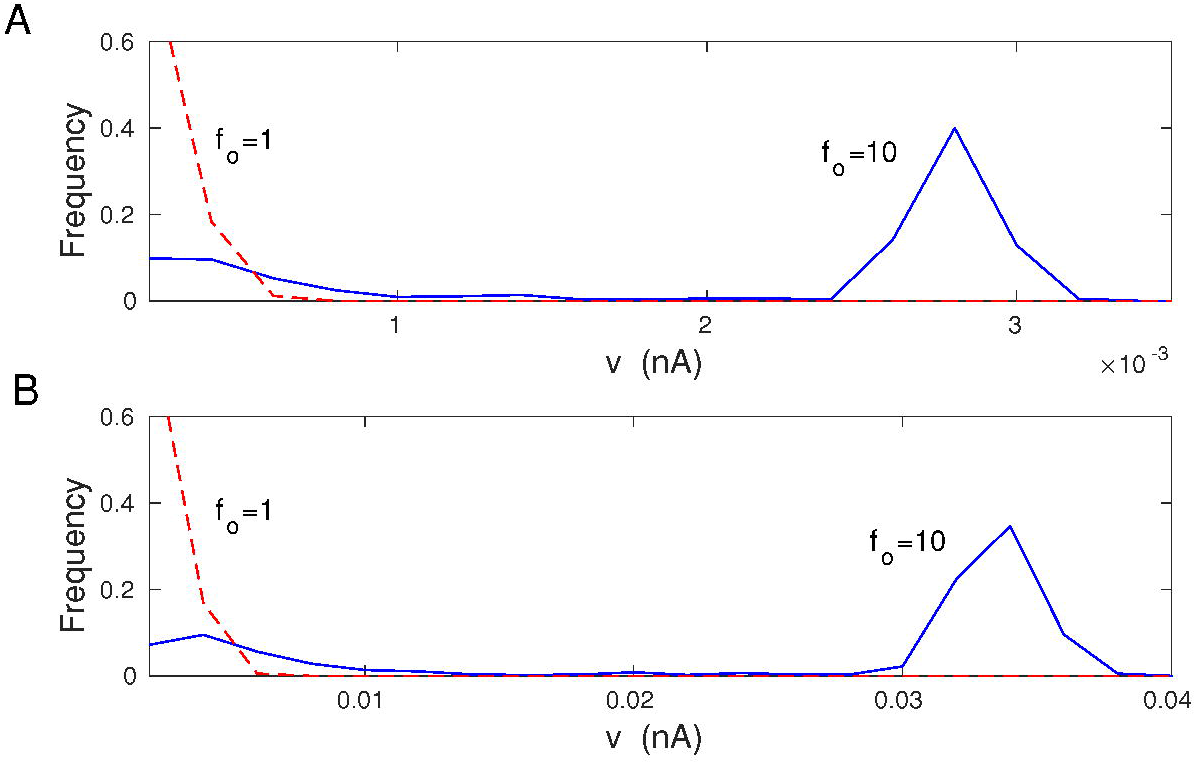
Distribution of synaptic current *v* in the stochastic mean-field model. Numerical solution of Eq. (3) and (6) for *σ_v_* > 0. In the monostable regime (*f_o_* = 1 Hz) the distribution is unimodal with a sharp peak around *v_d_* ≈ 0. In the bistable regime (*f_o_* = 10 Hz) the distribution is bimodal with two maxima around two fixed points *v_d_* and *v_u_*. (A) Parameters used: λ = 9 · 10^−7^ and *κ* = 0.001. (B) Parameters used: λ = 10^−5^ and *κ* = 0.012. In both (A) and (B) *σ_w_* = 0.1 nS.

Stationary average synaptic current 〈*v*〉, which is a measure of synaptic weights, increases weakly with presynaptic firing rate mean *f_o_* and its standard deviation *σ_f_* (Figs. 8 and 9). Mean-field values of 〈*v*〉 (computed from Eq. (41)) start to deviate from the exact numerical values (computed from Eqs. 1–2) for larger levels of synaptic noise *σ_w_* and for higher *f_o_* (Figs. 8 and 9).

**Fig. 8.**
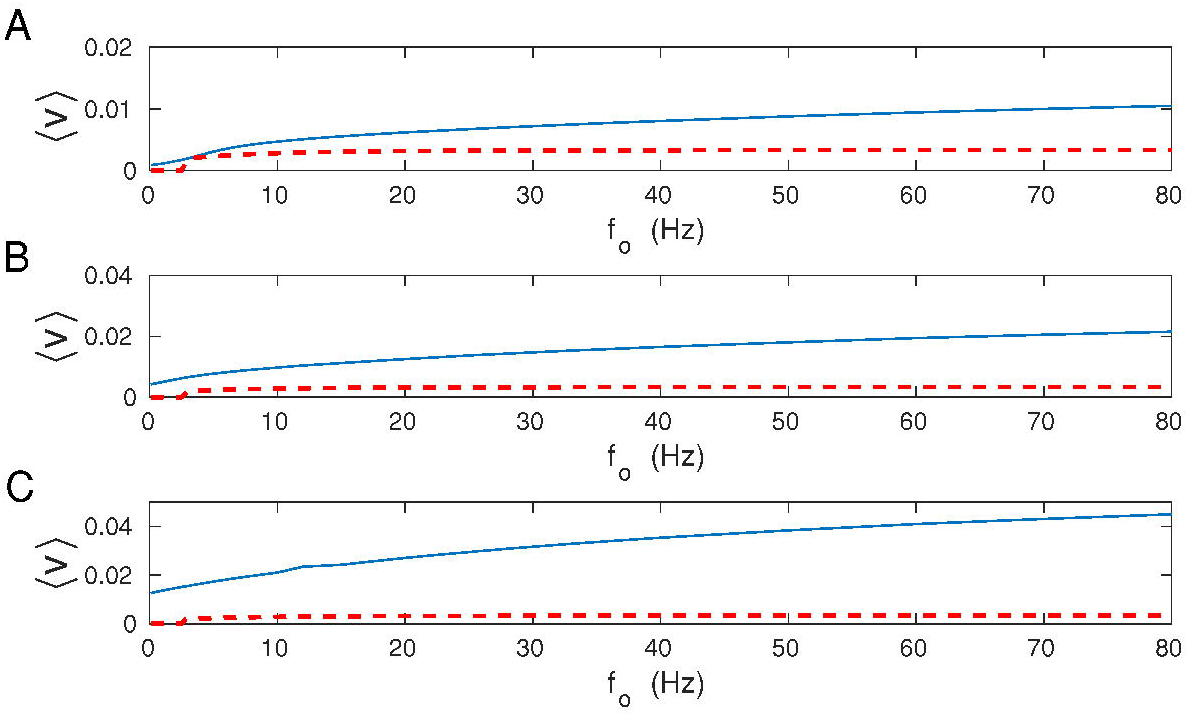
Average synaptic current 〈*v*〉 as a function of mean presynaptic firing rate: comparison of exact results with mean-field. (A) Dependence of 〈*v*〉 on *f_o_* for *σ_w_* = 0.02. (B) The same for *σ_w_* = 0.1. (C) The same for *σ_w_* = 0.5. For all panels solid lines correspond to exact result (from Eqs. 1–2), dashed lines to mean-field (from Eq. 44). The results are for λ = 9 · 10^−7^ and *κ* = 0.001.

**Fig. 9.**
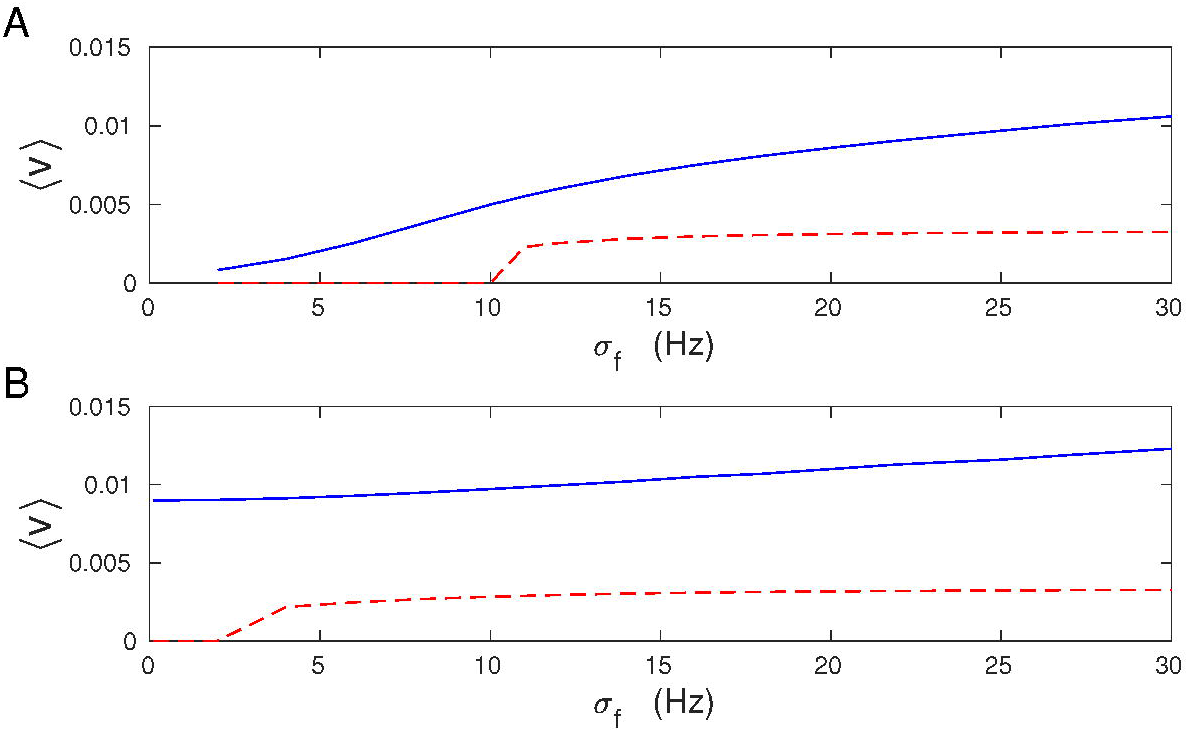
Average synaptic current 〈*v*〉 as a function of standard deviation of the presynaptic firing rate: comparison of exact results with mean-field. (A) Dependence of 〈*v*〉 on σf for *f_o_* =1 Hz. (B) The same for *f_o_* = 10 Hz. For all panels solid lines correspond to exact numerical result (from Eqs. 1–2), dashed lines to mean-field (from Eq. 44). The results are for λ = 9 · 10^−7^, *κ* = 0.001, and *σ_w_* = 0.1.

Stochastic Eq. (3) for the dynamics of v can be mapped into an equation for the dynamics of the probability distribution of v conditioned on *f_o_*, i.e. *P*(*v|f_o_*), described by a Fokker-Planck equation (see Eqs. (31–34) in the Methods). In the stochastic stationary state, characterized by the stationary probability distribution *P_s_* (*v|f_o_*), we can define a new and important quantity called an effective potential Φ(*v|f_o_*), which is a function of the synaptic current *v*. The effective potential Φ is proportional to the amount of energy associated with the synaptic plasticity described by Eq. (3), and it is equal to the integral of the right hand side of Eq. (3) with *σ_v_* = 0 (Van Kampen 2007; see Eq. (36) in the Methods). The explicit form of the effective potential Φ is

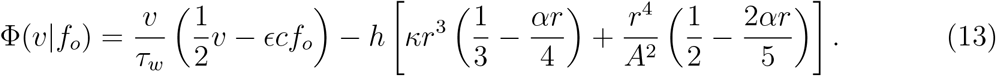

Note that the second term in Φ (with the large bracket) is proportional to the plasticity amplitude λ through *h*. This term depends on *v* through the firing rate *r* (see Eq. 6). In general, the functional form of the potential Φ(*v|f_o_*) determines the thermodynamics of synaptic memory, and thus it is an important function.

The shape of the potential Φ(*v|f_o_*) depends on the relative magnitude of the driving-plasticity *h* and the inverse of the decay time constant 1/*τ_w_* (Fig. 10A). In fact, there are two competing terms in Φ that are controlled by 1/*τ_w_* and h. The first term (~ 1/*τ_w_*) maintains monostability, while the second (~ *h*) promotes bistability. For *h* greater than the critical value *h_cr_* (Eq. 8), there is bistability and Φ has two minima at *v_u_* and *v_d_*, corresponding to up (strong) and down (weak) synaptic states (Fig. 10A), similar to the result for the deterministic limit. For very large *h*, there is again only one minimum related to strong synapses (Fig. 10A). The two minima are separated by a maximum corresponding to a potential barrier at *v_max_*. Metastable values of *v*, i.e. the minima and maximum of the potential, can be found from the condition *d*Φ/*dv* = 0, which is equivalent to finding the fixed points of Eq. (3) in the deterministic limit.

**Fig. 10.**
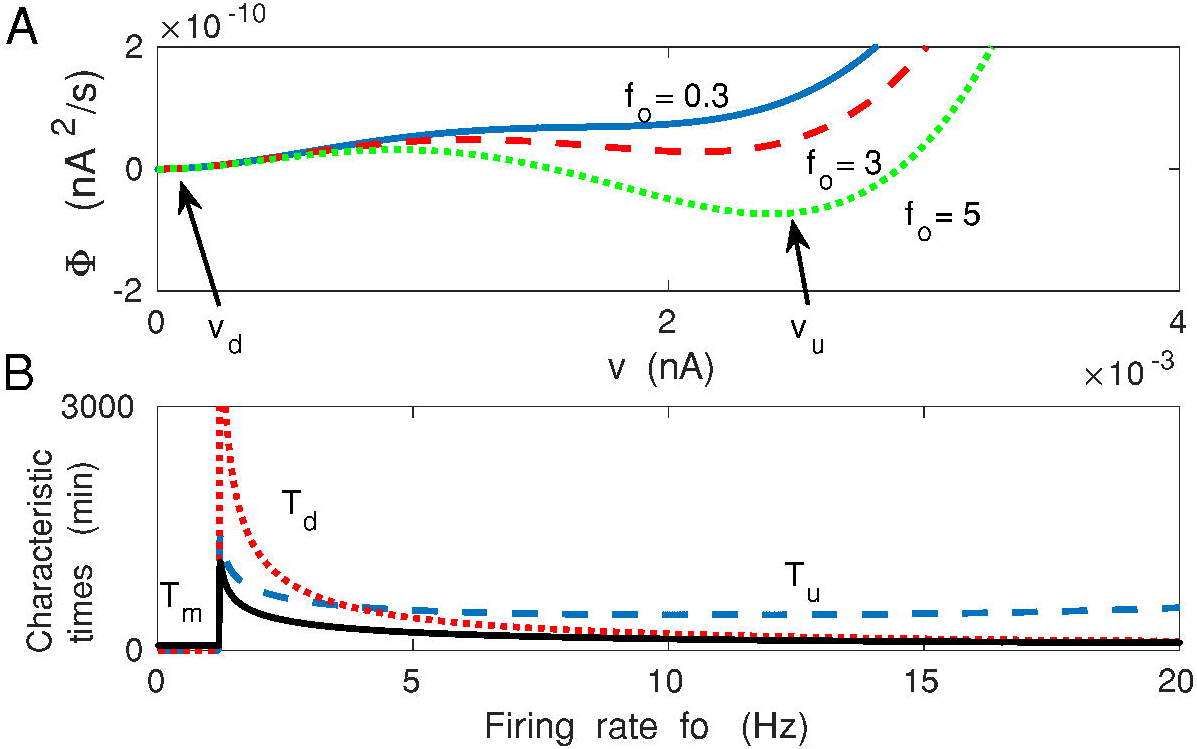
Effective synaptic potential, metastability, and memory lifetime: theory. (A) The metastable synaptic states can be described in probabilistic terms and correspond to minima of an effective potential Φ(*v|f_o_*). For weak presynaptic driving input *f_o_* the potential Φ has only one minimum at *v_d_* ~ *O*(*ϵ*), related to weak synapses. If *f_o_* is above a certain threshold, then the potential displays two minima, corresponding to bistable coexistence of weak and strong synapses (*v_d_* and *v_u_*). In the bistable regime, the synapses can jump between weak and strong states due to fluctuations in the input and/or synaptic noise. (B) Characteristic long times in the up (*T_u_*), down (*T_d_*) synaptic states, and memory lifetime as functions of presynaptic firing *f_o_*. Curves in (A) and (B) are for λ = 9 · 10^−7^ and *κ* = 0.001.

If we use a mechanical analogy and treat *v* as a spatial coordinate, then synaptic plasticity can be visualized as a movement in *v* space (state transitions), which is constrained by the energy related to Φ. This means that the shape of the function Φ(*v*) determines what kind of motions in v-space (state space) are possible or more likely. In particular, the binary nature of synaptic plasticity given by Eq. (3) can be described as transitions between two wells of the effective potential Φ(*v|f*_0_), corresponding to weak and strong synapses, or down and up synaptic states (e.g. Billings and van Rossum 2009; Graupner and Brunel 2012). These transitions, caused by intrinsic synaptic noise (*σ_w_*) and fluctuations in the presynaptic input (*σ_f_*), can be thought as a “hill climbing” process in the *v* space, which requires energy due to a barrier separating the two wells (Fig. 10). The dwelling times in both states (*T_u_,T_d_*) can be found from the classic Kramers “escape” formula (Eq. 47; Van Kampen 2007), and they are generally much larger than the time constant *τ_w_* (Fig. 10B).

We define the memory time *T_m_* of the synaptic system as a characteristic time needed to relax synaptic weights to their stationary values following a brief perturbation, or single memory event. Mathematically, it is equivalent to finding a relaxation time of the probability distribution *P*(*v|f_o_*) to its steady state distribution *P_s_*(*v|f_o_*) after a brief perturbation; see Eq. (50) in the Methods. The characteristic memory time *T_m_* is strictly related to the dwelling times *T_u_* and *T_d_* by Eq. (51), and they mutual relationship is depicted in Fig. 10B. Generally, the memory lifetime *T_m_* is very small in the monostable regime (*T_m_ ~ τ_w_*), i.e. for small presynaptic firing. However, it jumps by several orders of magnitude when synapses become bistable (i.e. when *h* ≈ *h_cr_*), but then *T_m_* monotonically decreases with increasing *f_o_* (Fig. 10B).

### Energy rate of synaptic plasticity

In this section we determine the energy rate, or metabolic rate, associated with stochastic BCM type synaptic plasticity. In a nutshell, energy is provided to the synaptic system to drive the plasticity related transitions between different synaptic weights associated mainly with the increase of synaptic weights. In the steady state this energy rate balances the energy dissipated due to the synaptic noise, which tends to decrease synaptic weights.

The plasticity related energy rate is determined both numerically, for the whole system of *N* synapses described by Eqs. (1–2), and analytically for the mean-field approximation described by a single Eq. (3). The numerical procedure for the whole system is described in the Methods (see section “Numerical simulations of the full synaptic system”), and the analytical results are described below.

The power dissipated by synaptic plasticity *Ė* in the mean-field approximation is proportional to the average temporal rate of the effective potential decrease, i.e. − 〈*d*Φ(*v|f*_0_)/*dt*〉, where 〈…〉 denotes averaging with respect to the probability distribution *P*(*v|f_o_*). Since the potential Φ(*v|f*_0_) depends on time only through *v*, after rearranging we get 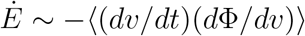. Thermodynamically, this formula is equivalent to the entropy production rate associated with the stochastic process described by Eq. (3), and represented by the effective potential Φ(*v|f_o_*) (Nicolis and Prigogine 1977; Lan et al 2012, Mehta and Schwab 2012; Tome 2006; Tome and de Oliveira 2010). The synaptic plasticity energy rate per synapse 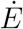 can be found analytically using 1/*N* expansion, and in the steady state takes the form (see Methods):

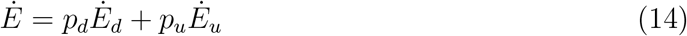

where *Ė_d_* and *Ė_u_* are the energy rates dissipated, respectively, in the down and up synaptic states, which have the occupancies *p_d_* and *p_u_*. The energy rates *Ė_d_* and *Ė_u_* are given by

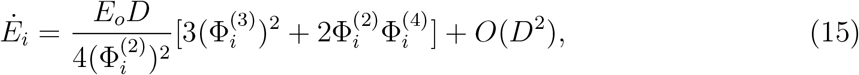

where *i* = *d* (down state) or *i* = *u* (up state). The symbols Φ_*i*_ and 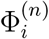 denote values of the potential Φ(*v*) and its *n*-th derivative with respect to *v* for *v* = *v_i_*. The symbol *E_o_* is the characteristic energy scale for variability in synaptic (spine) conductance, and it provides a link with underlying molecular processes (see the Methods). For convenience, we defined a new noise related parameter *D*, which is

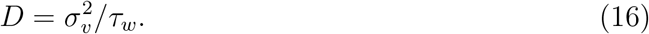

*D* can be viewed as the effective noise amplitude, and *D* relates to the number of synapses *N* as *D* ~ 1/*N*.

Note that in Eq. (15) the terms of the order *O*(1) disappear, and the first nonzero contribution to *Ė* is of the order *O*(1/*N*), since *D* ~ 1/*N*. Moreover, to have nonzero power in this order, the potential Φ(*v*) must contain at least a cubic nonlinearity.

Eqs. (14) and (15) indicate that energy is needed for plasticity processes associated with the potential Φ “hill climbing”, which is in analogy to the energy needed for a particle trapped in a potential well (of a certain shape) to escape. The energetics of such a “motion” in the *v*-space depends on the shape of the potential, which is mathematically accounted for by various higher-order derivatives of Φ. Thus, a fraction of synapses that were initially in the down state can move up the potential gradient to the up state by overcoming a potential barrier, but this requires the energy that is proportional to 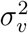 and to the derivatives of the potential. By analogy, a similar picture holds for synapses that were initially in the up state. The prefactor D in Eq. (15) indicates that the transitions up↔down, as well as local fluctuations near these states, cost energy that is proportional to the intrinsic synaptic noise (~ *σ_w_*) and presynaptic activity (including its fluctuations *f_o_* and *σ_f_*). The important point is that if there is no intrinsic spine noise (*σ_w_* = 0), then there are no transitions between the up and down states in the steady state, and consequently there is no energy dissipation (*σ_v_* = 0), regardless of the fast presynaptic input magnitude. Likewise for very long decay of synaptic weights, *τ_w_* ↦ ∞, corresponding to very slow synaptic plasticity (and the lack of the decay term in Eq. 1), there is no energy used. In such a noiseless stationary state, the plasticity processes described by Eq. (3) are energetically costless, since there are no net forces that can change synaptic weight, or mathematically speaking, that can push synapses in the v-space. (This is not true under non-stationary conditions when there is some temporal variability in one or more parameters in Eq. 3, leading to dissipation, but the focus here is on the steady state). This situation resembles the so-called “fluctuation-dissipation” theorem known from statistical physics (Nicolis and Prigogine 1977; Van Kampen 2007; Risken 1996), where thermal fluctuations always cause energy loss. In our case, this fluctuation-dissipation relationship underlines a key role of thermodynamic fluctuations for the metabolic load of synaptic plasticity.

We can compare the energy rate coming from the mean-field (Eq. 3) to the energy rate computed numerically for the full synaptic system described by Eqs. (1–2). The results are presented in Figs. 11 and 12. Generally, a better agreement between mean-field and exact results is achieved for intermediate synaptic noise *σ_w_* and also for intermediate values of mean presynaptic firing rate *f_o_*. For larger *σ_w_*, in the regions close to mono-bistability transitions, there are peaks in the mean-field *Ė* that are absent in the numerical *Ė* (Fig. 11). These peaks are the artifacts of the approximation methods used in the mean-field. Moreover, Fig. 11 shows that the energy rate *Ė* mostly increases steadily with *f_o_* (the exact result). The exception is a narrow interval near the mono-to bi-stability regions, where *Ė* slightly decreases (Fig. 11). The energy rate also steady increases with the standard deviation in the presynaptic firing *σ_f_* (Fig. 12).

**Fig. 11.**
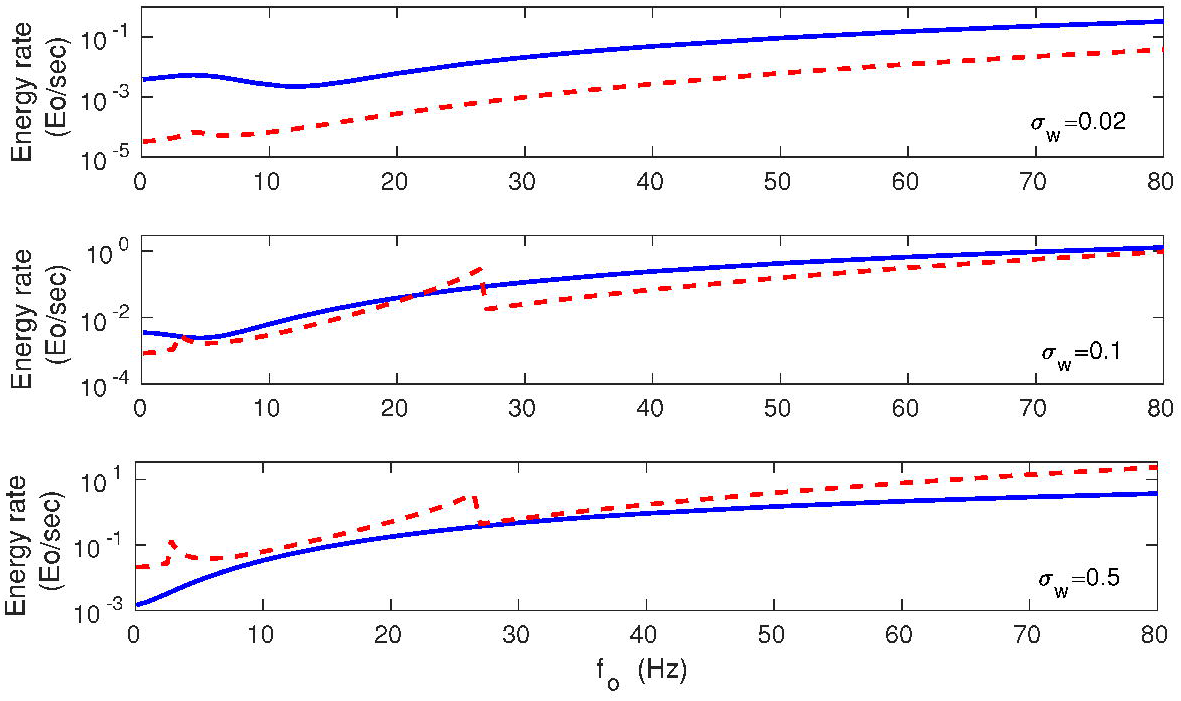
Energy rate *Ė* as a function of mean presynaptic firing rate: comparison of exact results with mean-field. Results are for *σ_w_* = 0.02 nS (upper panel), *σ_w_* = 0.1 nS (middle panel), and *σ_w_* = 0.5 nS (lower panel). Solid lines correspond to exact numerical results for the whole system of *N* synapses obtained from Eq. (79). Dashed lines correspond to the mean-field approach (Eqs. 14–15). The best agreement between exact and mean-field results is for the intermediate *σ_w_*, and not too small *f_o_*. Note two peaks in the mean-field result for *Ė* (corresponding to mono ↔ bistability transitions) for larger noise *σ_w_*, which are absent in the exact results. These peaks are the artifacts of the approximation method in the mean-field. All plots are for λ = 9 · 10^−7^, *κ* = 0.001, and *σ_f_* = 10 Hz.

**Fig. 12.**
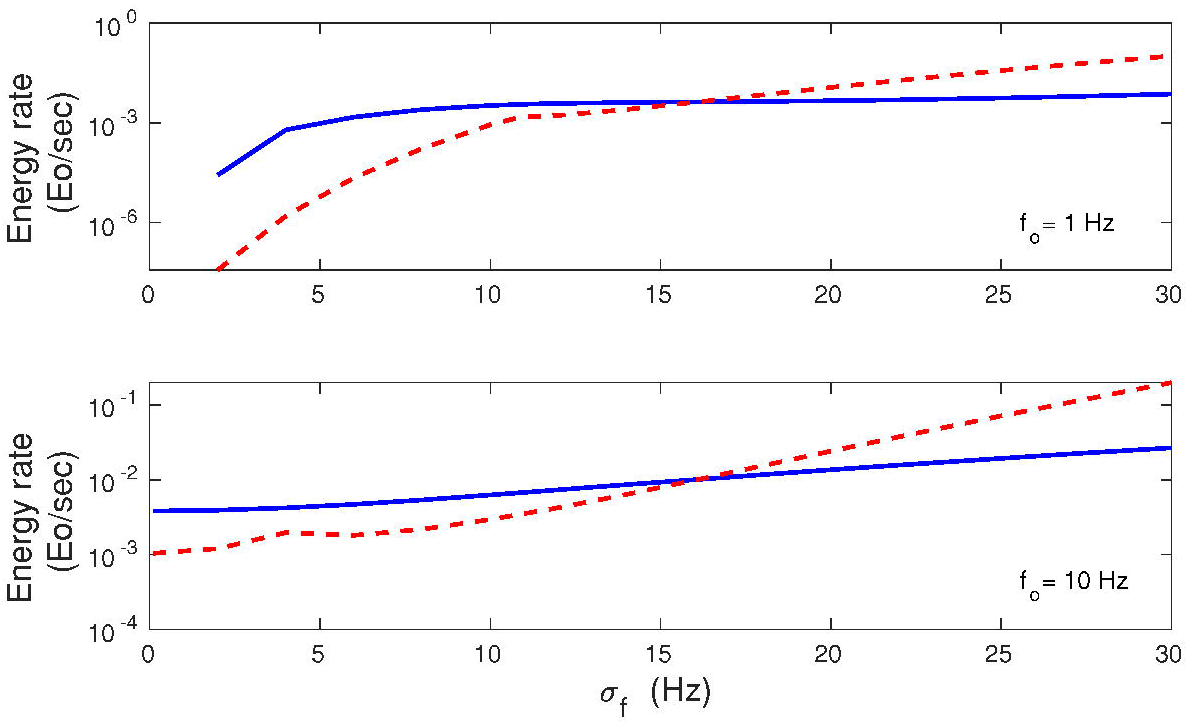
Energy rate *Ė* as a function of standard deviation of the presynaptic firing rate: comparison of exact results with mean-field. Results are for *f_o_* = 1 Hz (upper panel), and *f_o_* = 10 Hz (lower panel). Solid lines correspond to exact numerical results obtained from Eq. (79), and dashed lines correspond to the mean-field results obtained from Eqs. (14–15). A better agreement between exact and mean-field results is obtained for the higher *f_o_*. All plots are for λ = 9 · 10^−7^, *κ* = 0.001, and *σ_w_* = 0.1 nS.

A next interesting question is how the plasticity energy rate depends on the synaptic weights? In Fig. 13 we plot the dependence of *Ė* on the average synaptic current (*v*), which is proportional to the synaptic weights and spine size (Kasai et al 2003). It is clear that the synaptic energy rate related to plasticity grows nonlinearly with 〈*v*〉. For small 〈*v*〉, the energy rate *Ė* depends weakly on 〈*v*〉, whereas for large 〈*v*〉 it increases strongly with 〈*v*〉 (Fig. 13).

**Fig. 13.**
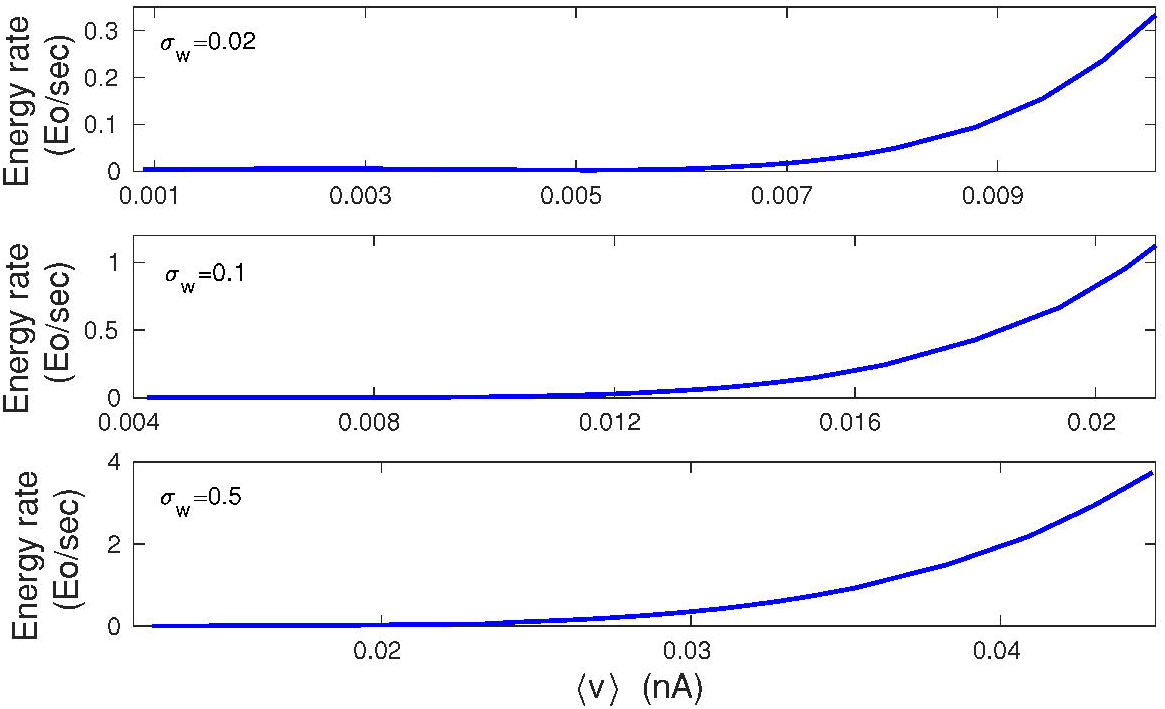
Energy rate *Ė* as a function of average synaptic current 〈*v*〉: exact numerical results. Note a sharp increase in E for larger 〈*v*〉. Energy rate is calculated from Eq. (79), and 〈*v*〉 is calculated from Eqs. (1–2). All plots are for λ = 9 · 10^−7^, *κ* = 0.001, and *σ_f_* = 10 Hz.

Which dependence of *Ė* is stronger: on *f_o_* or on 〈*v*〉? In Fig. 14 it is shown that the energy rate *Ė* increases nonlinearly both with *f_o_* and with 〈*v*〉, but the dependence on the average synaptic current 〈*v*〉 is much steeper.

**Fig. 14.**
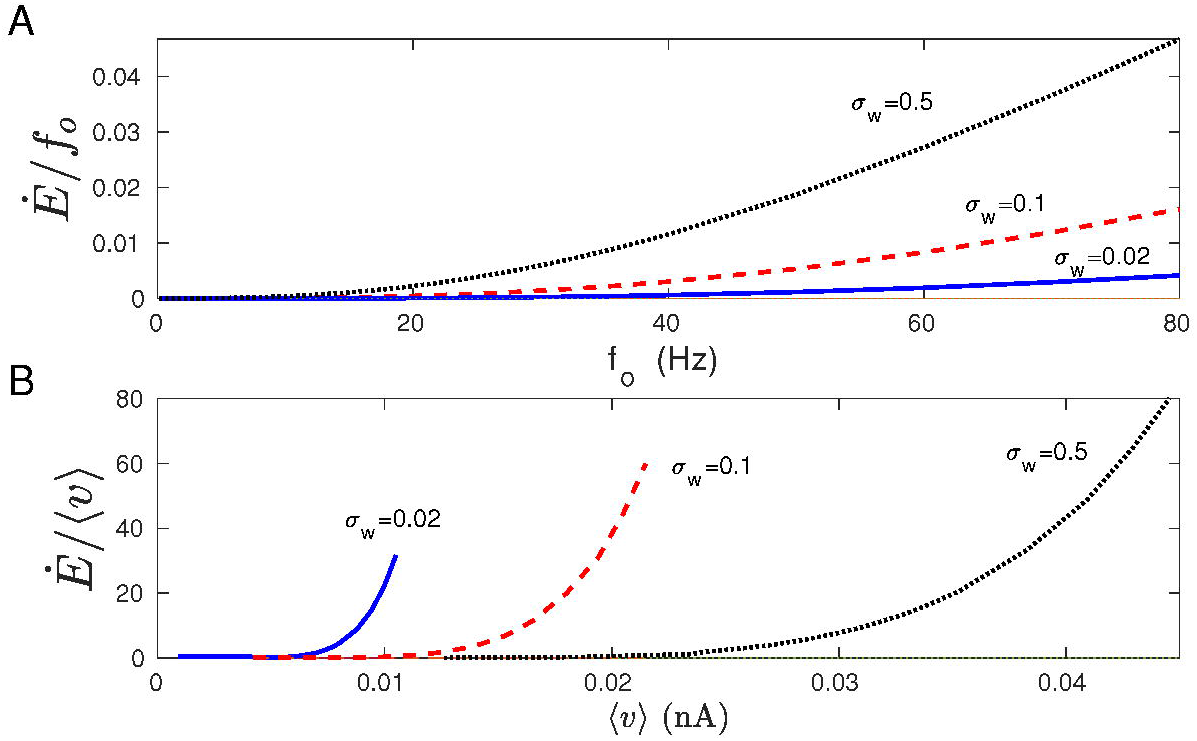
Nonlinear dependence of energy rate on presynaptic firing rate and average synaptic current: exact numerical results. (A) The ratio *Ė/f_o_* as a function of *f_o_*. (B) The ratio *Ė*/〈*v*〉 as a function of 〈*v*〉. The energy rate IE depends nonlinearly both on *f_o_* and v, but the dependence on the synaptic current 〈*v*〉 is more steep. Energy rate is calculated from Eq. (79), and 〈*v*〉 is calculated from Eqs. (1–2). All plots are for λ = 9 · 10^−7^, *κ* = 0.001, and σf = 10 Hz.

### Energy cost of plastic synapses as a fraction of neuronal electric energy cost: comparison to experimental data

In order to assess the magnitude of the synaptic plasticity energy rate, we compare it to the rate of energy consumption by a typical cortical neuron for its electric activities related to fast synaptic transmission, action potentials and maintenance of the resting potential (Attwell and Laughlin 2001). The neural spiking activity and synaptic transmission are known to consume the majority of the neural energy budget (Attwell and Laughlin 2001; Harris et al 2012; Karbowski 2012). The ratio of the total energy rate used by plastic synapses *NĖ* to the neuron’s energy rate *Ė_n_* (given by Eq. (63) in the Methods) is computed for different presynaptic firing rates *f_o_*, various levels of synaptic noise *σ_w_*, and for different cortical regions. The results for macaque and human cerebral cortex are shown in Figs. 15 and 16. These plots indicate that the synaptic plasticity contribution depends strongly on the level of synaptic noise *σ_w_*; the higher the noise the larger the ratio *NĖ/Ė_n_*. Higher firing rates *f_o_* also tend to increase that ratio but not that strongly, and the dependence is nonmonotonic (Figs. 15 and 16). Generally, the value of *NĖ/Ė_n_* ranges from negligible (~ 10^−4^ − 10^−3^) for small/intermediate noise (*σ_w_* = 0.1 nS), to substantial (~ 10^−2^ − 10^0^) for very large noise (*σ_w_* = 2 nS). The results are qualitatively similar across different cortical regions within one species, as well as between human and macaque cortex, despite large differences in the cortical sizes of both species (Figs 15 and 16). Small quantitative differences are the result of small differences in the synaptic densities between areas and species.

**Fig. 15.**
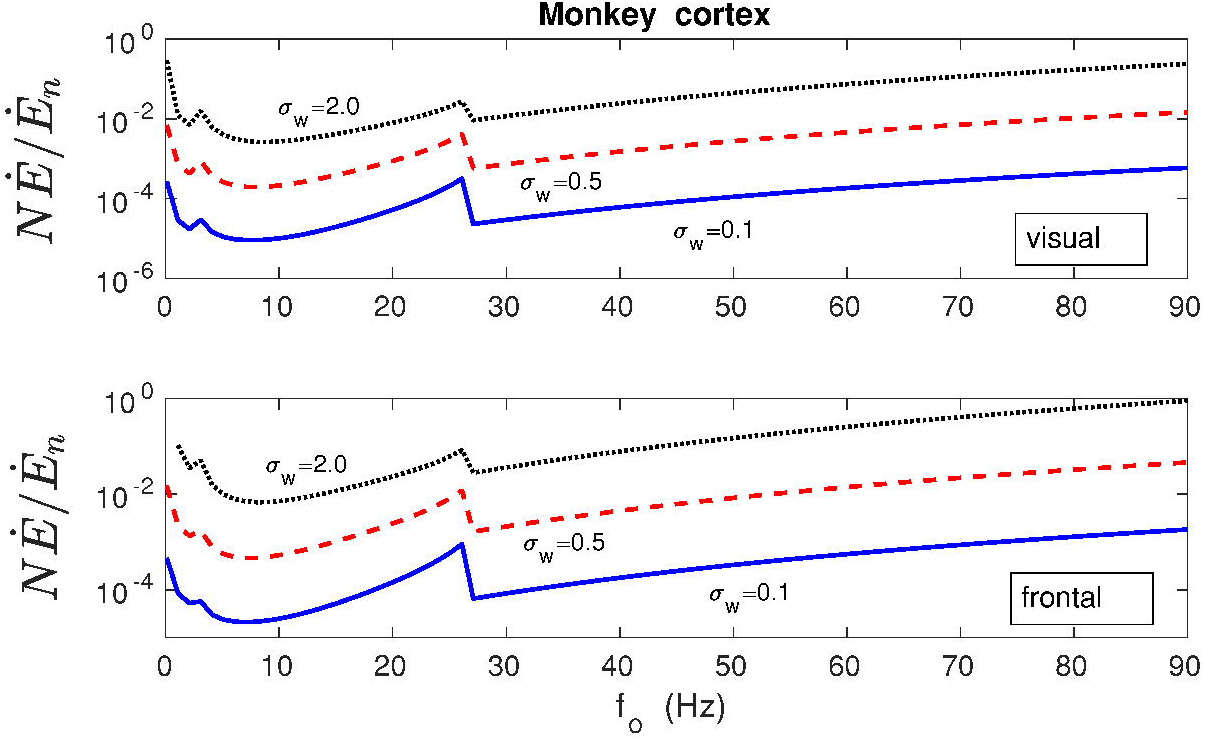
Energy cost of synaptic plasticity as a fraction of neuron’s electric energy cost in the cerebral cortex of macaque monkey. The ratio of the total energy rate used by plastic synapses *NĖ* (chemical energy) to neuron’s energy rate *Ė_n_* (electric energy used mainly for fast synaptic transmission and neural spiking) as a function of presynaptic firing rate *f_o_* for different levels of synaptic noise *σ_w_*, and different regions of the macaque cortex (visual and frontal). Note that the energy contribution of plastic synapses to the neuron’s energy budget depends strongly on the synaptic noise level. For weak and intermediate noise this contribution is mostly marginal. For very large noise (*σ_w_* = 2 nS) it can be substantial, but only for very large firing rates. The neuron’s energy rate *Ė_n_* was computed using Eq. (63), while the plasticity energy rate of all synapses *NĖ* was computed from Eqs. (14–15). All plots are for λ = 9 · 10^−7^, *κ* = 0.001, and *σ_f_* = 10 Hz.

**Fig. 16.**
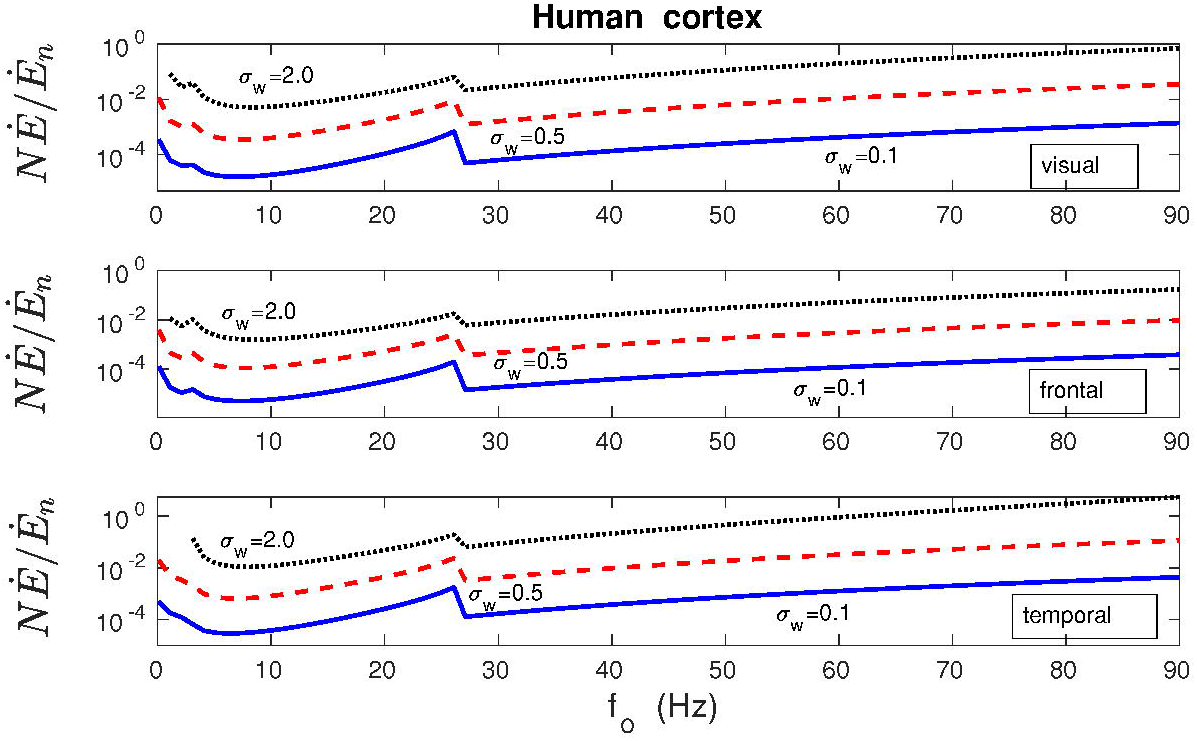
Similar as in Fig. 15, but for human cerebral cortex. Overall, the ratio of the energy rate of plastic synapses to the neuron’s electric energy rate for human cortex is very similar to the one for macaque cortex.

### Information encoded by plastic synapses

In our model, information or memory about the mean input *f_o_* is written in the population of synapses, represented by the synaptic current *v*. In the stochastic steady state, the synaptic current *v* is characterized by probability distribution *P_s_*(*v|f*_0_), which is related to the potential Φ(*v|f*_0_). This means that information encoded in synapses also depends on the structure of the potential (Eq. 13).

The accuracy of the encoded information can be characterized by Fisher information *I_F_* (Cover and Thomas 2006). In general, larger *I_F_* implies a higher coding precision. Fisher information, related to synaptic current *v*, can be derived analytically (see the Methods). In the limit of small effective noise amplitude *D* we obtain:

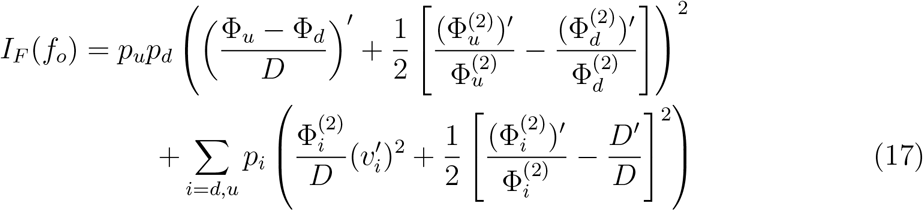

where *p_i_* denote the fractions of synapses in the up (*i = u*) and down (*i = d*) states, and the prime denotes a derivative with respect to *f_o_*. Note that the effective noise amplitude *D* depends on *f_o_*, since *σ_v_* depends on *f_o_* and 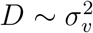 (see Eq. (5)).

The first term in Eq. (17) proportional to *p_d_p_u_* is of the order of ~ 1/*D*^2^ ~ *N*^2^, and it appears only in the bistable regime (both *p_d_* and *p_u_* must be nonzero). This term depends on the difference in the potentials between up and down states. The second term in Eq. (17) proportional to the weighted sums of *p_d_* and *p_u_* is of the order of ~ 1/*D* ~ *N*, and it is always present regardless of mono- or bistability. Thus, the first term is much bigger than the second for small *D*, which is the primary reason why *I_F_* (and coding accuracy) is several orders of magnitude larger when synapses are bistable (see below). Because Fisher information *I_F_*(*f_o_*) is either proportional to 1/*D*^2^ ~ *N*^2^ (in the bistable regime) or to 1/*D ~ N* (in the monostable regime), it implies that many synapses are much better in coding the presynaptic firing *f_o_* than a single synapse.

Equation for Fisher information (Eq. 17) indicates that there is no simple relationship between *I_F_* and the synaptic current *v*. Rather, *I_F_* depends in a nonlinear manner on the derivatives of synaptic currents in up and down states 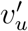 and 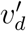. This follows form the fact that the potential Φ depends in a complicated way on *v* (see Fig. 10, and Eq. (13)).

### Accuracy and lifetime of synaptically stored information vs plasticity energy rate

How the long-term energy used by synapses relates to the accuracy and persistence of stored information? The above results indicate that *Ė* and *I_F_* depend inversely on the synaptic noise *σ_v_* (or *D*), suggesting that its lowering should be beneficial since gain in information is accompanied by a decrease in synaptic energy rate.

A more complicated picture emerges if other parameters are varied, notably driving presynaptic input *f_o_*, at different regimes of mono- and bistability (Fig. 17). At the onset of bistability, Fisher information *I_F_* and memory lifetime *T_m_* both increase dramatically, whereas the plasticity energy rate *Ė* increases mildly. Approximate meanfield calculations of *Ė* provide a small peak at the transition point, but more exact numerical calculations of *Ė* based on Eqs. (1–2) indicate a smooth behavior (with a slight decrease), which suggest that the small peak in the mean-field is an artifact of the approximation (Fig. 17). Taken together, this implies that a high improvement in information coding accuracy and its retention, in the initial region of bistability, do not involve a huge amounts of energy. On the contrary, the corresponding energy cost is rather small.

**Fig. 17.**
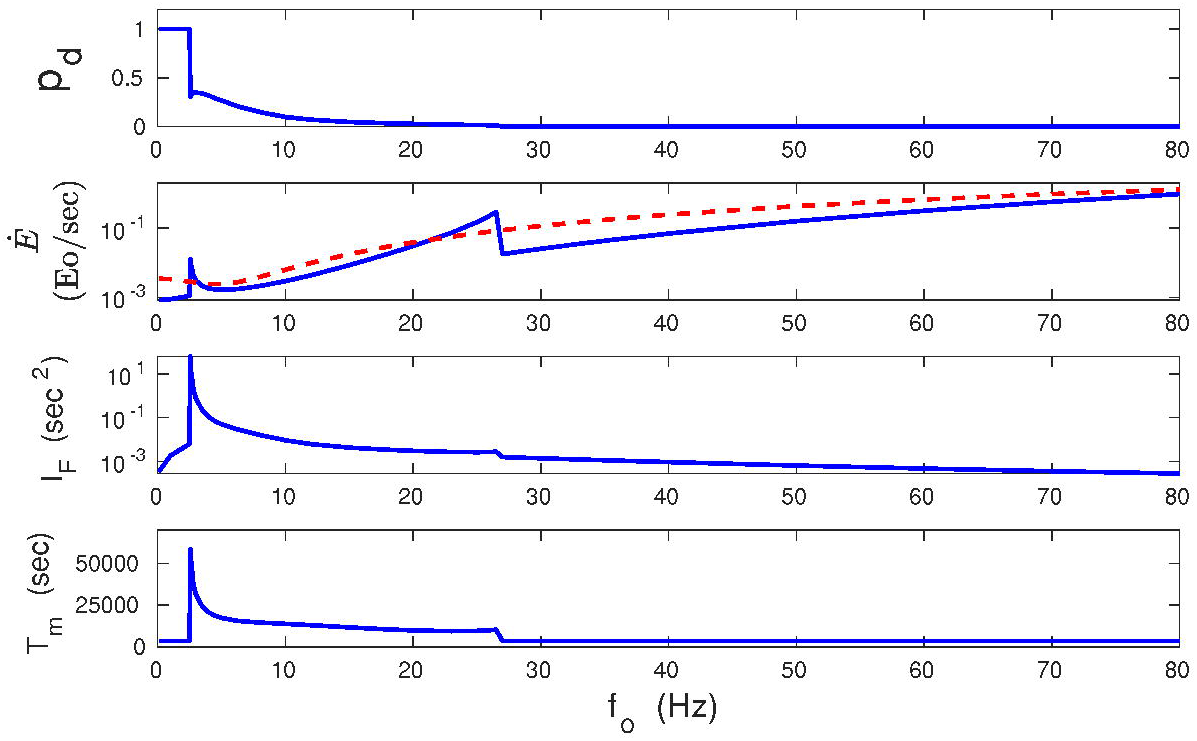
Comparison of synaptic plasticity energy rate with accuracy and lifetime of stored information as a function of presynaptic firing rate. Dependence of *p_d_, Ė, I_f_*, and on firing rate *f_o_*. Fisher information *I_F_* and memory lifetime *T_m_* have large peaks at the onset of bistability, whereas the synaptic energy rate *Ė* increases only mildly. Beyond the transition point to bistability, *I_F_* and *T_m_* exhibit a different dependence on *f_o_* than *Ė*. The former two quantities decrease while the latter increases with *f_o_*. In the dependence of *Ė* on *f_o_*, the solid line corresponds to the mean-field approximation, and the dashed line to the exact numerical result. All plots are for λ = 9 · 10^−7^, *κ* = 0.001, and *σ_w_* = 0.1 nS.

For higher *f_o_*, deeper in the bistability region, there is a different trend. In this coexistence region, *Ė* increases monotonically, while *I_F_* and *T_m_* decrease, which in turn indicates an inefficiency of information storing. However, even here the huge values in *I_F_* and *T_m_* overcome the growth in *Ė*. For even higher *f_o_*, in the monostable phase with strong synapses only, *Ė* still increases monotonically, whereas *I_F_* and *T_m_* further decrease to the levels similar for very small *f_o_*. Consequently, the biggest gains in synaptic information precision and lifetime per energy used (*I_F_/Ė* and *T_m_/Ė*) are achieved for the bistable phase only (Fig. 18). Interestingly, the gains in the information precision and lifetime depend nonmonotonically on the plasticity amplitude λ, and there are some optimal values of λ that are different for the gains *I_F_/Ė* and *T_m_/Ė* (Fig. 19).

**Fig. 18.**
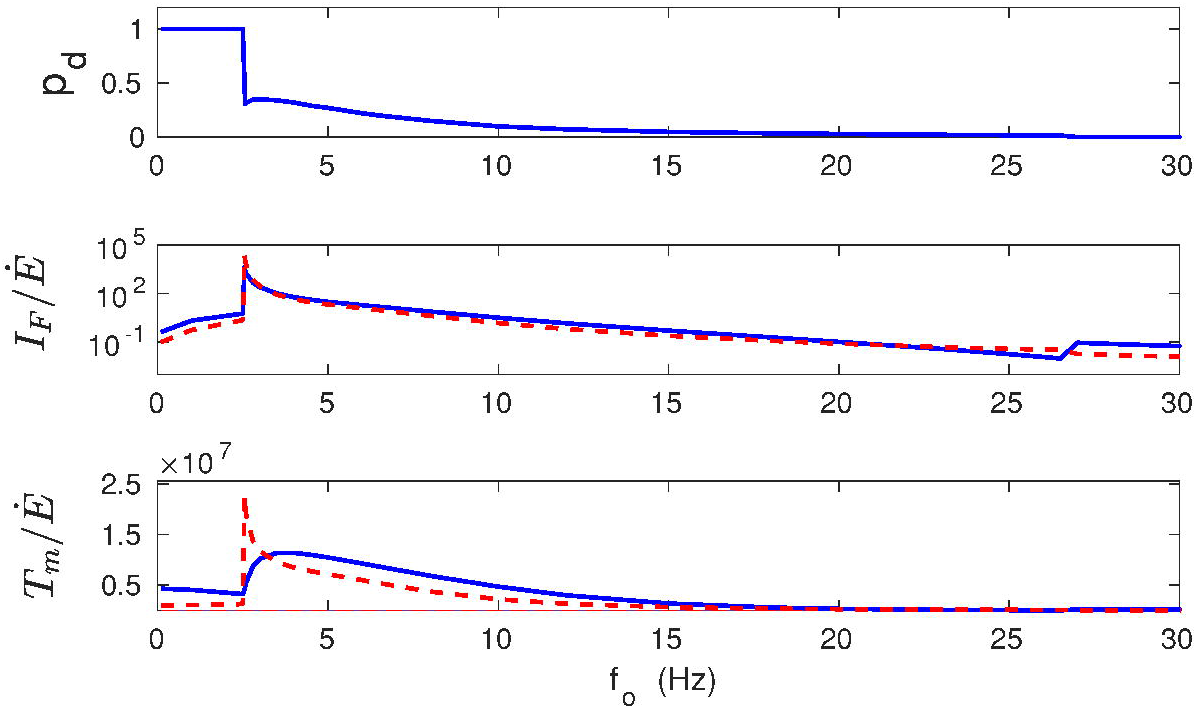
Gains in information accuracy and lifetime per synaptic energy used as functions of presynaptic firing rate. Fraction of synapses in the down state *p_d_* (upper panel), the ratio *I_F_/Ė* (middle panel) and *T_m_/Ė* (lower panel) as functions of *f_o_*. The biggest gains in information and memory lifetime are at the transition point to the bistability. Solid lines correspond to the energy rate calculated in the mean-field, and dashed lines to the energy rate calculated numerically. All plots are for λ = 9 · 10^−7^, *κ* = 0.001, and *σ_w_* = 0.1 nS.

**Fig. 19.**
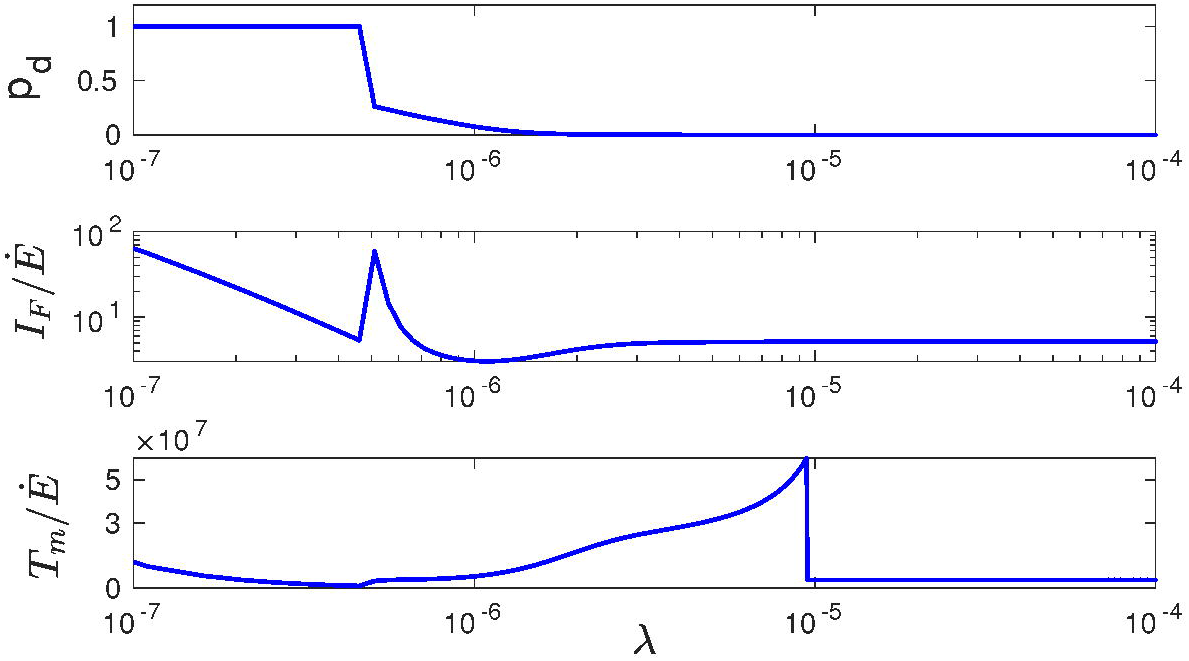
Gains in information accuracy and lifetime per synaptic energy used as functions of plasticity amplitude. Fraction of synapses in the down state *p_d_* (upper panel), the ratio *I_F_/Ė* (middle panel) and *T_m_/Ė* (lower panel) as functions of λ. Note the sharp peaks for *I_F_/Ė* at the transition point from mono-to bi-stability, and for *T_m_/Ė* at the transition point from bi-to mono-stability. Solid lines correspond to the energy rate calculated in the mean-field approach. All plots are for *f_o_* = 10 Hz, *κ* = 0.001, and *σ_w_* = 0.1 nS.

Taken together, these results suggest that storing of accurate information in synapses can be relatively cheap in the bistable regime, and thus metabolically efficient.

### Precision of coding memory is restricted by sensitivity of synaptic plasticity energy rate on the driving input

The above results suggest that synaptic energy utilization does not limit directly the coding precision of a stimulus, because there is no simple relationship between Fisher information and power dissipated by synapses. However, a careful inspection of the curves in Fig. 17 suggests that there might be a link between *I_F_* and the derivative of *Ė* with respect to the driving input *f_o_*. In fact, it can be shown that in the most interesting regime of synaptic bistability, in the limit of very weak effective noise *D* → 0, we have either (see the Methods)

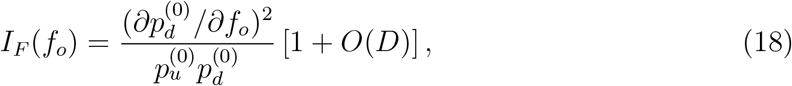

or equivalently

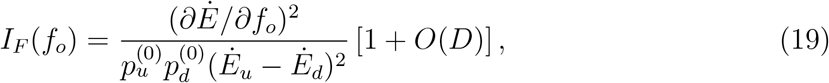

where 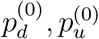 are the fractions of synapses in the down and up states (weak and strong synapses) in the limit *D* ↦ 0. It is important to stress that simple formulas (18) and (19) have a general character, since they do not depend explicitly on the potential Φ, and thus they are independent of the plasticity type model. Equation (18) shows that synaptic coding precision increases greatly for sharp transitions from mono-to bistability, since then 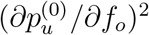 is large. Additionally, Eq. (19) makes an explicit connection between precision of synaptic information and nonequilibrium dissipation. Specifically, the latter formula implies that to attain a high fidelity of stored information, the energy used by synapses *Ė* does not have to be large, but instead it must change sufficiently quickly in response to changes in the presynaptic input.

We can also estimate a relative error ef in synaptic coding of the average presynaptic firing *f_o_*. This error is related to Fisher information by a Cramer-Rao inequality 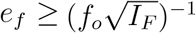 (Cover and Thomas 2006). Using Eq. (19), in our case this relation implies

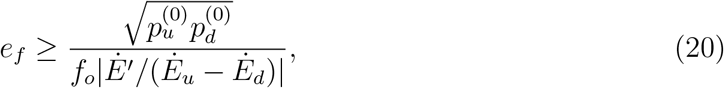

where the prime denotes the derivative with respect to *f_o_*. The value of the product *p_u_p_d_* is in the range from 0 to 1/4. In the worst case scenario for coding precision, i.e. for 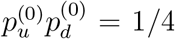, this implies that a 10% coding error (*e_f_* = 0.1), corresponds to the relative sensitivity of the plasticity energy rate on presynaptic firing *f*_0_1*Ė*’/(*Ė_u_—Ė_d_* | = 5. Generally, the larger the latter value, the higher precision of synaptic coding. In our particular case, this high level of synaptic coding fidelity is achieved right after the appearance of bistability (Fig. 17).

## 3. Discussion

### Summary of the main results

In this study, the energy cost of long-term synaptic plasticity was determined and compared to the accuracy and lifetime of an information stored at excitatory synapses. The main results of this study are:

a. Formulation of the dynamic mean-field of the extended BCM synaptic plasticity model (Eqs. 3–5).
b. Energy rate of plastic synapses increases nonlinearly both with the presynaptic firing rate (Figs. 11 and 14) and with average synaptic current or weights (Figs. 13 and 14).
c. Coding of more accurate information in synapses need not require a large energy cost (cheap long-term information). The accuracy of stored information about presynaptic input can increase by several orders of magnitude with only a mild increase in the plasticity energy rate at the onset of bistability (Figs. 17 and 18).
d. The accuracy of information stored at synapses and its lifetime are not limited by the available energy rate, but by the sensitivity of the energy rate on the presynaptic firing. For very weak synaptic noise the coding accuracy at plastic synapses (Fisher information) is proportional to the square of derivative of the plasticity energy rate with respect to mean presynaptic firing (Eq. 19).
e. Energy rate of synaptic plasticity, which is of chemical origin, constitutes in most cases only a tiny fraction of neuron’s energy rate associated with fast synaptic transmission and action potentials, which are of electric origin. That fraction can be substantial only for very large synaptic noise and presynaptic firing rates (Figs. 15 and 16).

### Discussion of the main results

The dynamic mean-field for synaptic plasticity was derived analytically by applying (i) the timescale separation between neural and synaptic plasticity activities, and (ii) dimensional reduction of the original synaptic system. The formulated mean-field of the synaptic current *v* seems to work reasonably well for average 〈*v*〉 if intrinsic synaptic noise *σ_w_* is small and presynaptic firing rates *f_o_* are not too high (Fig. 8). For larger *σ_w_* and *f_o_*, the mean-field value 〈*v*〉 diverges form the exact numerical average calculated from Eqs. (1–2).

The mean-field approximation to the synaptic energy rate *Ė* was additionally derived in the limit of small effective noise D (either large *N* or small *σ_w_*, or both). Surprisingly, the mean-field approximation for *Ė* works better for intermediate noise *σ_w_* than for its smaller values (Fig. 11). For those intermediate values of *σ_w_*, the energy rate *Ė* calculated in the mean-field is close to that calculated numerically in the whole neurophysiological interval of *f_o_* variability (Fig. 11, middle panel). It seems that the primary reason for the breakdown of the mean-field for 〈*v*〉 and *Ė* is the way the integrals in Eqs. (43) and (57) were approximated. In those integrals, it was assumed that *x*_*d*1_ (the lower limit of integration) tends to —∞ for *D* ↦ 0, which however is not always true, since *x*_*d*1_ ~ *v_d_* ~ *ϵ*, and *v_d_* can be very small for very small value of e (see Eq. 11), especially if *f_o_* is small. As a consequence, the real value of *x*_*d*1_ can be in the range —(0.1 − 1), even for very small *D*.

Comparing the mean-field *Ė* to its numerical values suggests that the peaks in the mean-field approximation of *Ė* are artifacts (Fig. 11). They are the result of some (small) differences in the exact location of the transitions points mono/bistability between mean-field and the numerics. This causes certain errors in the relative magnitudes of *p_d_* and *p_u_*, which leads to over- or under-estimates in the mean-field values of *Ė* (Eq. 14).

Nonlinear increase of plasticity energy rate with presynaptic firing rate *f_o_* and average synaptic current *v* (Figs. 11, 13, and 14) suggests that high presynaptic activities and large synapses/spines are metabolically costly (there exists a positive correlation between synaptic current and size; see Kasai et al (2003)). Consequently, it seems that large firing rates and synaptic weights (proportional to average synaptic current) should not be preferred in real neural circuits. This simple conclusion is qualitatively in line with experimental data for cortical neurons, showing low mean firing rates and weak mean synaptic weights, with skewed distributions (Buzsaki and Mizuseki 2014).

The most striking result of this study is that precise memory storing about presynaptic firing rate *f_o_* does not have to be metabolically expensive. Strictly speaking, the information encoded at synapses, i.e., its accuracy and lifetime, do not have to correlate positively with the energy used by synapses (Fig. 17). Such a correlation is only present at the onset of synaptic bistability, where a large increase in information precision (*I_F_*) and lifetime (*T_m_*) is accompanied only by a mild increase in the energy rate. This suggests an energetic efficiency of stored information in the bistable synaptic regime, i.e., relatively high information gain per energy used (Fig. 18). Moreover, the results in Fig. 18 show that there exists an optimal value of the presynaptic firing rate for which the information gain per energy, as well as memory lifetime per energy, are maximal. An additional support for the metabolic efficiency of synaptic information comes from the fact that energy used *Ė* and coding precision *I_F_* depend the opposite way on the effective noise amplitude *D* (compare Eqs. (15) and (17)), and thus, *I_F_* increases, while *Ė* decreases with decreasing *D*. Because *D* ~ 1/*N*, this also implies that *I_F_ ~ N*^2^ in the bistable regime, i.e., that more synapses (large *N*) are much better at precise coding of mean presynaptic firing than a single synapse (*N* = 1). Taken together, these findings are compatible with a study by Still et al (2012) showing that abstract stochastic systems with memory, operating far from thermodynamic equilibrium, can be the most predictive about an environment if they use minimal energy.

Estimating an external variable is never perfect, and it is shown here that synaptic coding accuracy (*I_F_*) relates to the derivative of the energy rate with respect to an average input. The fundamental relationship linking memory precision and synaptic metabolic sensitivity is present in Eq. (19), which is valid regardless of the specific plasticity mechanism, as long as synapses can exist in two metastable states, in the limit of very small synaptic noise *D*. This binary synaptic nature is a key feature enabling a high fidelity of long-term synaptic information (Petersen et al 1998), despite ongoing neural activity, which is generally detrimental to information storing (Fusi et al 2005). Specifically, for realistic neurophysiological parameters, it is seen from Fig. (17) that the relative coding error in synapses 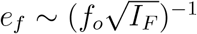 can be as small as 0.03 − 0.1 (or 3 − 10%) near the onset of bistability. However, away from that point the error gets larger. Thus, again it seems that there exist an optimal firing rate *f_o_* for which coding accuracy is maximal and quite high, despite large fluctuations in presynaptic neural activities (large *σ_f_* in relation to *f_o_*).

Neural computation is thought to be metabolically expensive (Aiello and Wheeler 1995; Laughlin et al 1998; Attwell and Laughlin 2001; Karbowski 2007 and 2009; Niven and Laughlin 2008; Harris et al 2012), and it must be supported by cerebral blood flow and constrained by underlying microvasculature and neuroanatomy (Karbowski 2014 and 2015). It is shown here that an important aspect of this computation, namely long-term synaptic plasticity involved in learning and memory, constitutes in most cases only a small fraction of that neuronal energy cost associated mostly with fast synaptic transmission and spiking (Figs. 15 and 16). Specifically, for intermediate/large synaptic noise (*σ_w_* = 0.1 and 0.5 nS), metabolic cost of synaptic plasticity can maximally be on a level of 1 − 10% of the electric neuronal cost, both for human and macaque monkey (Figs. 15 and 16). Higher levels of synaptic plasticity cost (maximally 100% of electric cost) are possible, but only for very large synaptic noise, *σ_w_* = 2.0 nS (Figs. 15 and 16). The latter value is, however, unlikely because it is 20 times larger than the mean values of synaptic weights *w_i_* (see Fig. 2B), and thus, it seems that higher costs of synaptic plasticity are physiologically implausible. Taken together, these results suggest that a precise memory storing can be relatively cheap, which agrees with empirical estimates presented in (Karbowski 2019).

### Discussion of other aspects of the plasticity model

In this study, an extended BCM model of synaptic plasticity is introduced and solved. There are 3 additional elements in our model (Eq. 1) that are absent in the classical BCM plasticity rule: weight decay term (~ 1/*τ_w_*), synaptic noise 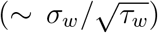, and the nonlinear dependence of the postsynaptic firing rate on synaptic input (Eq. 6). Moreover, it is assumed here that presynaptic firing rates fluctuate stochastically and fast around a common mean *f_o_* with standard deviation *σ_f_*. These features make the behavior of our model significantly different from the behavior of the classical BCM model (Bienenstock et al 1982). In particular, due to the stochasticity of synaptic weights, our model does not exhibit an input selectivity, in contrast to the classical BCM rule. Input selectivity in the classical BCM means that the largest static presynaptic firing rate “selects” its corresponding synapse by increasing its weight, in such a way that the weights of all other synapses decay to zero. In our model this never happens, because all synapses are driven on average by the same input, and more importantly, synaptic noise constantly brings all synapses up and down in an unpredictable fashion. For these reasons, the mean-field approach proposed here, although mathematically correct, does not make sense for the classical BCM rule (no weights decay, no noise) if our goal is studying input selectivity, because in that model only one synapse is effectively present at the steady-state, and there is no need for large *N* approach.

The main reasons for choosing the mean-field approach, and constructing a single dynamical equation for the population averaged synaptic current *v* are: (i) we wanted to treat analytically the multidimensional stochastic model given by Eqs. (1–2), and (ii) the variable *v* emerges as a natural choice, since *r* in Eqs. (1–2) depends only on one variable, precisely on *v* (see Eq. 6). The feature (ii) makes Eqs. (3) and (6) a closed mathematical system of just two equations that can be handled analytically.Another practical reason behind introducing the dynamic mean-field is that it enables us to obtain explicit formulae for synaptic plasticity energy rate and Fisher information (coding accuracy).

In deriving the dynamic mean-field we assumed that the time constant related to *w_i_* dynamics, i.e. *τ_w_*, is much larger than the time constant related to the sliding threshold *θ*, which is *τ_θ_*. This is in agreement with empirical observations and estimations, since *τ_w_* must be of the order of 1 hr to be consistent with slice experiments, showing wiping out synaptic potentiation after about 1 hr when presynaptic firing becomes zero (Frey and Morris 1997; Zenke et al 2013). (Note that *τ_w_* refers to the decay of synaptic weights to the baseline value ea, and it should not be confused with a characteristic time of plasticity induction, which is controlled by the product λ*f_i_r* in Eq. (1) and which can be much faster, ~ minutes (Petersen et al 1998; O’Connor et al 2005).) On the other hand, the time constant *τ_θ_* must be smaller than about 3 min for stability reasons (Zenke and Gerstner 2017; Zenke et al 2013), and it even has been estimated to be as small as ~ 12 sec (Jedlicka et al 2015).

Although individual synapses in the original model Eqs. (1–2) exhibit bistability (see Figs. 1 and 2), this bistability has a collective character. That is, if most synaptic weights are initially weak, then they all converge into a lower fixed point. On the other hand, if a sufficient fraction of synaptic weights is initially strong, then they all converge into an upper fixed point (Fig. 1). This means that the majority of synapses participate in a coordinated switching between up and down states, due to effective noise (internal and external). This mechanism is probably different from the mechanism found in Petersen et al (1998) and O’Connor et al (2005), where bistability was reported on a level of a single synapse, independent of other synapses. (However, from these papers it is difficult to judge how long the potentiation lasts in the absence of presynaptic stimulation). Our scenario for bistability is conceptually closer to the model of synaptic bistability proposed by Zenke et al 2015 (Zenke et al 2015), which also emerges on a population level. Interestingly, both models, the one presented here and the one in (Zenke et al 2015), exhibit the so-called anti-Hebbian plasticity, in the sense that LTP (i.e. *v* > 0) appears for low firing rates, instead of LTD as for classical BCM rule. However, in the present model the initial LTP window is very narrow, and appears for very small postsynaptic firing rates *r* < (*cf*_0_/*κ*)*ϵ* ~ *O*(*ϵ*). This feature is necessary for stable bistability, and does not contradict experimental results on BCM rule verification (Kirkwood et al 1996), showing LTD for low firing rates. The reason is that these experiments were performed for firing rates above 0.1 Hz, leaving uncertainty about LTP vs. LTD for very low activity levels (or very long times).

The cooperativity in synaptic bistable plasticity found here is to some extent similar to the data showing that neighboring dendritic spines interact and tend to cluster as either strong or weak synapses (Govindarajan et al 2006; Govindarajanet al 2011). These clusters can be as long as single dendritic segments, which is called “clustered plasticity hypothesis” (Govindarajan et al 2006; Govindarajan et al 2011). However, the difference is that in the present model there are no dendritic segments, and spatial dependence is averaged over, which leads effectively to one synaptic “cluster” either with up or down states.

### Metabolic cost of synaptic plasticity in the mean-field: intuitive picture

The formula for the plasticity energy rate (Eqs. 15) contains various derivatives of the effective potential Φ, which encodes the plasticity rules for synaptic weights. In this scenario, the synaptic plasticity corresponds to a driven stochastic motion of the population averaged postsynaptic current *v* in the space constrained by the potential Φ, in analogy to a ball moving on a rugged landscape with a ball coordinate corresponding to v. Because our potential can exhibit two minima separated by a potential barrier, the plasticity considered here can be viewed as a stochastic process of “hill climbing”, or transitions between the two minima (the idea of “synaptic potential” was used also in van Rossum et al (2000), Billings and van Rossum (2009), Graupner and Brunel (2012)).

The energy rate of plastic synapses *Ė* (or power dissipated by plasticity) is the energy used for climbing the potential shape in v-space, and it is proportional to the average temporal rate of decrease in the potential, − 〈*d*Φ/*dt*〉, due to variability in *v*. In terms of thermodynamics, the plasticity energy rate *Ė* is equivalent to the entropy production rate, because synapses like all biological systems operate out of thermodynamic equilibrium with their environment and act as dissipative structures (Nicolis and Prigogine 1977). Dissipation requires a permanent influx of energy from the outside (provided by blood flow, see e.g. Karbowski (2014)) to maintain synaptic structure, which in our case is the distribution of synaptic weight. A physical reason for the energy dissipation in synapses in the steady state is the presence of noise (both internal synaptic ~ *σ_w_*, and external presynaptic ~ *σ_f_*), causing fluctuations, that tend to wipe out the pattern of synaptic weights. Thermodynamically speaking, this means reducing the synaptic order and thus increasing synaptic entropy. To preserve the order, this increased entropy has to be “pumped out”, in the form of heat, by investing some energy in the process, which relates to ATP consumption.

### Thermodynamics of memory storing and bistability

The general lack of high energetic demands for sustaining accurate synaptic memory may seem non-intuitive, given an intimate relation between energy and information known from classical physics (Leff and Rex 1990). For example, transmitting 1 bit of information through synapses is rather expensive and costs 10^4^ ATP molecules (Laughlin et al 1998), and a comparative number of glucose molecules (Karbowski 2012), which energetically is much higher (~ 10^5^*kT*) than a thermodynamic minimum set by the Landauer limit (~ 1*kT*) (Landauer 1961). Additionally, there are classic and recent theoretical results that show dissipation-error tradeoff for biomolecular processes, i.e., that higher coding accuracy needs more energy (Lan et al 2012; Mehta and Schwab 2012; Barato and Seifert 2015; Bennett 1979; Lang et al 2014). How can we understand our result in that light?

First, there is a difference between transmitting information and storing it, primarily in their time scales, and faster processes generally need more power (see also below). Second, it is known from thermodynamics that erasing an information can be more energy costly than storing information (Landauer 1961; Bennett 1982), since the former process is irreversible and is always associated with energy dissipation, and the latter can in principle be performed very slowly (i.e. in equilibrium with the environment)without any heat released. In our system, the information is maximal for intermediate presynaptic input generating metastability with two synaptic states (Fig. 2). If we decrease the input below a certain critical value, or increase it above a certain high level, our system becomes monostable, which implies that it does not store much information (entropy is close to zero). Thus, the transition from bistability to monostability is equivalent to erasing the information stored in synapses, which according to the Landauer principle (Landauer 1961; Berut et al 2012) should cost energy.

Third, the papers showing energy-error tradeoff in biomolecular systems (Lan et al 2012; Mehta and Schwab 2012; Lang et al 2014; Bennett 1979; Barato and Seifert 2015) use fairly linear (or weakly nonlinear) models, while in our model the plasticity dynamics is highly nonlinear (see Eqs. 1, 3, and 6). Additionally, we consider the prediction of an external variable (average input *f_o_*), in contrast to some of the biomolecular models (Bennett 1979; Barato and Seifert 2015), which dealt with estimating errors in an internal variable.

### Cost of synaptic plasticity in relation to other neural costs

The energy cost of synaptic plasticity is a new and an additional contribution to the overall neural energy budget considered before and associated with fast signaling (action potentials, synaptic transmission, maintenance of negative resting potential), and slow nonsignaling factors (Attwell and Laughlin 2001; Engl and Attwell 2015). The important distinction between slow synaptic plasticity dynamics and fast signaling is that the former is of chemical origin (protein/receptor interactions), while the latter is of electric origin (ionic movement against gradients of potential and concentration).Consequently these two, although coupled but to a large extent, separate phenomena have different characteristic time and energy scales, which results in rather small energy cost of synaptic plasticity in relation to the fast electric cost (Figs. 15 and 16).

The earlier studies of the neuronal energy cost (Attwell and Laughlin 2001; Engl and Attwell 2015) provided important order of magnitude estimates based on ATP turnover rates, but they had mainly a phenomenological character and cannot be directly applied to nonlinear dynamics underlying synaptic plasticity. Contrary, the current approach and the complementary approach taken in (Karbowski 2019) are based on “first principles” taken from non-equilibrium statistical physics and in combination with neural modeling can serve as a basis for future more sophisticated calculations of energy used in excitatory synapses, possibly with inclusion of some molecular detail (e.g. Lisman et al 2012; Miller et al 2005; Kandel et al 2014).

The calculations performed here indicate that the energy dissipated by synaptic plasticity increases nonlinearly with presynaptic firing rate (Fig. 11). The dependence on presynaptic firing is consistent with a strong dependence of CaMKII autophosphorylation level on Ca^2+^ influx frequency to a dendritic spine (De Koninck and Schulman 1998), which should translate to a similar dependence of ATP consumption rate related to protein activation on presynaptic firing. Moreover, these results raise the possibility of observing or measuring the energetics of synaptic plasticity for high firing rates. It is hard to propose a specific imaging technique for detecting enhanced synaptic plasticity, but nevertheless, it seems that techniques relying on spectroscopy, e.g., near-infrared spectroscopy with its high spatial and temporal resolution, could be of help.Regardless of whether the energetics of synaptic plasticity is observable or not, it could have some functional implications. For example, it was reported that small regional decreases in glucose metabolic rate associated with age, and presumably with synaptic decline, lead to significant cognitive impairment associated with learning (Gage et al 1984).

A relatively small cost of plasticity in relation to neuronal cost of fast electric signaling (Figs. 15 and 16) is in some part due to relatively slow dynamics of spine conductance decay, quantified by *τ_w_* ~ 1 hr (Frey and Morris 1997; Zenke et al 2013), since *Ė* ~ 1/*τ_w_* in Eqs. 15 and 16. The time scale *τ_w_* characterizes the duration of early LTP on a single synapse level. On a synaptic population level, characterized by synaptic current v, the duration of early LTP is given by *T_m_* (memory maintenance of a brief synaptic event), which can be of the order of several hours.

Late phases of LTP and LTD, during which memory is consolidated, are much slower than *τ_w_* and they are governed by longer timescales of the orders of days/weeks (Ziegler et al 2015; Redondo and Morris 2011). Consequently, one can expect that such plasticity processes, as well as equally slow homeostatic synaptic scaling (Turrigiano and Nelson 2004), should be energetically inexpensive. Nevertheless, there are experimental studies related to long-term memory cost in fruitfly that claim that memory in general is metabolically costly (Mery and Kawecki 2005; Placais and Preat 2013; Placais et al 2017). However, the problem with those papers is that they do not measure directly the energy cost related to plasticity in synapses, but instead they estimate the global fly metabolism, which indeed affects long-term memory (Mery and Kawecki 2005; Placais and Preat 2013). In a recent paper by Placais et al (2017) it was found that upregulated energy metabolism in dopaminergic neurons is correlated with long-term memory formation. However, again, no measurement was made directly in synapses, and thus it is difficult to say how much of this enhanced neural metabolism can be attributed to plasticity processes and how much to enhanced neural and synaptic electric signaling (spiking and transmission). It is important to stress that the energy cost of protein synthesis, process believed to be associated with long-term memory consolidation (Kandel et al 2014), was estimated to be very small, on a level of ~ 0.03 − 0.1% of the metabolic cost of fast synaptic electric signaling related to synaptic transmission (Karbowski 2019; see also below for an alternative estimate). Consequently, it is possible that memory induction, maintenance, and consolidation involve a significant increase in neural activity and hence metabolism, but it seems that the majority of this energy enhancement goes for upregulating neural electric activity, not for chemical changes in plastic synapses.

The energetics of very slow processes associated with memory consolidation were not included in the budget of the energy scale *E_o_* (present in Eq. 15, and estimated in the Methods), since we were concerned only with the early phases of LTP and LTD, which are believed to be described by BCM model (both standard and extended). Nevertheless, for the sake of completeness, we can estimate the energy cost of the late LTP and LTD, as well as energy requirement of mechanical changing of spine volume (also not included in the budget of *E_o_*).

Protein synthesis, which is associated with l-LTP and l-LTD, underlines synaptic consolidation and scaling (Kandel et al 2014). There are roughly 10^4^ proteins in PSD including their copies (Sheng and Hoogenraad 2007), on average each with ~ 400 − 500 amino acids, which are bound by peptide bonds. These bonds require 4 ATP molecules to form (Engl and Attwell 2015), which is 4 · 20*kT* of energy (Phillips et al 2012). This means that chemical energy associated with PSD proteins is about (3.2 − 4.0) · 10^8^kT, i.e. (1.6 − 2.0) · 10^7^ ATP molecules, or equivalently (1.4 − 1.75) · 10^−12^ J. Given that an average lifetime of PSD proteins is 3.7 days (Cohen et al 2013), we obtain the energy rate of protein turnover as ~ (4.6 − 5.8) · 10 ^−18^ W, or 52 − 65 ATP/s per spine. For human cerebral cortex with a volume of 680 cm^3^ (Hofman 1988) and average density of synapses 3 · 10^11^ cm^−3^ (Huttenlocher and Dabholkar 1997), we have 2 · 10^14^ synapses. This means that the global energy cost of protein turnover in spines of the human cortex is (9.2 − 11.5) · 10^−4^ W, or equivalently (1 − 1.3) · 10^16^ ATP/s, which is extremely small (~ 0.01%) as human cortex uses about 5.7 Watts of energy (Karbowski 2009).

The changes in spine volume are related directly to the underlying dynamics of actin cytoskeleton (Honkura et al 2008; Cingolani and Goda 2008). We can estimate the energy cost of spine size using a mechanistic argument. Dendritic spine grows due to pressure exerted on the dendrite membrane by actin molecules. The reported membrane tension is in the range (10^−4^ − 1) kT/nm^2^ (Phillips et al 2009) with the upper bound being likely an overestimate, given that it is close to the so-called rapture tension (1 − 2 kT/nm^2^), when the membrane breaks (Phillips et al 2009). A more reasonable value of the membrane tension seems to be 0.02 kT/nm^2^, as it was measured directly (Stachowiak et al 2013). Taking this value, we get that to create a typical 1 *μ*m^2^ of stable spine requires 2 · 10^4^*kT* or 10^3^ ATP molecules. Since the actin turnover rate in spine is 1/40 sec^−1^ (Honkura et al 2008), which is also the rate of spine volume dynamics, we obtain that the cost of maintaining spine size is 25 ATP/s. This value is comparable but two-fold smaller than the ATP rate used for PSD protein turnover per spine (52 − 65 ATP/s) given above.

How do the costs of protein turnover and spine mechanical stability relate to the energy cost of e-LTP and e-LTD calculated in this paper using the extended BCM model? From Fig. 11, we get that the latter type of synaptic plasticity uses energy in the range (10^−3^ − 10^0^)*E_o_* (solid lines for exact numerical results) per second per spine, depending mainly on firing rate and synaptic noise. Since the energy scale *E_o_* = 2.3 · 10^4^ ATP (see the Methods), we obtain that the energy cost of the plasticity related to e-LTP and e-LTD is 23 − 23000 ATP/s, i.e., its upper range can be 400 times larger than the contributions from protein turnover and spine volume changes. This result strongly suggests that the calculations of the energetics of synaptic plasticity based on the extended BCM model provide a large portion of the total energy required for the induction and maintenance of synaptic plasticity.

## 4. METHODS

### Neuron model

We consider a sensory neuron with a nonlinear firing rate curve (so called class one, valid for most biophysical models) and with activity adaptation given by (Ermentrout 1998; Ermentrout and Terman 2010)

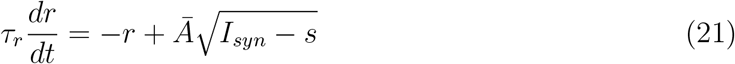

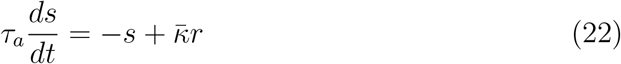

where *r* is the instantaneous neuron firing rate with mean amplitude *Ā*, s is the adaptation current (or equivalently self-inhibition) with the intensity 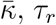 and *τ_a_* are the time constants for variability in neural firing and adaptation, and *I_syn_* is the total excitatory synaptic current to the neuron provided by *N* excitatory synapses, i.e., *I_syn_* ~ ∑_*i*_ *f_i_w_i_*. If *I_syn_* < *s* in Eq. (21), then this equation simplifies and becomes *τ_r_dr/dt* = − *r*. In order to ensure a saturation of the firing rate *r* for very large number of synapses *N*, and for *s* to be relevant in this limit, *Ā* and 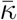 must scale as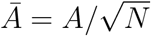 and 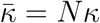. In a mature brain *N* can fluctuate due to structural plasticity, but we assume in agreement with the data (Sherwood et al 2020; DeFelipe et al 2002) that there is some well defined average value of *N*.

We assume that the neuron is driven by stochastic presynaptic firing rates *f_i_* (*i* = 1,…,*N*) that change on a much faster time scale *τ_f_* than the synaptic weights *w_i_*. Additionally, we assume that the fast variability in presynaptic firing rates is stationary in a stochastic sense, i.e., the probability distribution of *f_i_* does not change in time. Consequently, for each time step *t*, in the stationary stochastic state we can write

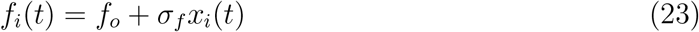

where *f_o_* is the mean firing rate of all presynaptic neurons and *σ_f_* denotes the standard deviation in the variability of *f_i_*. The variable *x_i_* is the Gaussian random variable, which reflects noise in the presynaptic neuronal activity. For the noise *x_i_* we have the following averages (Van Kampen 2007): 〈*x_i_*〉_x_ = 0 and 〈*x_i_x_j_*〉_*x*_ = *δ_ij_*, where the last equality means that different *x_i_* are independent, which also implies that fluctuations in different firing rates *f_i_* are statistically independent. Eq. (23) allows negative values of *f_i_*, which is not realistic. However, in analytical calculations this is not a problem, because we use only average of *f_i_* and its standard deviation *σ_f_*. In numerical simulations, we prevent the negative values of *f_i_* by setting *f_i_* = 0, whenever *f_i_* becomes negative.

Given Eq. (23), one can easily verify the following average

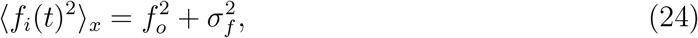

Equation (24) indicates that presynaptic firing rates fluctuate around average value *f_o_* with standard deviation *σ_f_*. The important point is that these fluctuations are fast, on the order of *τ_f_* (~ 0.1 − 1 sec), which is much faster than the timescale *τ_w_*. Eq. (24) is also used below.

### Definition of synaptic current per spine *v*

The synaptic current *I_syn_* has two additive components related to AMPA and NMDA receptors, *I_syn_* = *I_ampa_ + I_nmda_*, with the receptor currents

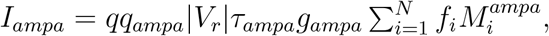

and

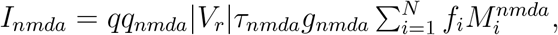

where *q* is the probability of neurotransmitter release, *V_r_* is resting membrane potential of the neuron (we used the fact that the reversal potential for AMPA/NMDA is close to 0 mV; Ermentrout and Terman (2010)), *g_ampa_* and *g_nmda_* are single channel conductances of AMPA and NMDA receptors, *q_ampa_* and *q_nmda_* are probabilities of their opening with characteristic times *τ_ampa_* and *τ_nmda_*. The symbols 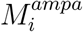 and 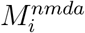 denote AMPA and NMDA receptor numbers for spine *i*. Data indicate that during synaptic plasticity the most profound changes are in the number of AMPA receptors *M^ampa^* and opening probability of NMDA *q_nmda_* (Kasai et al 2003; Huganir and Nicoll 2013; Matsuzaki et al 2004). We define the excitatory synaptic weight *w_i_* as a weighted average of AMPA and NMDA conductances, i.e.,

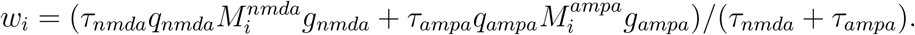

This enables us to write the synaptic current per spine, i.e. *v* = *I_syn_/N* (which is more convenient to use than *I_syn_*), as

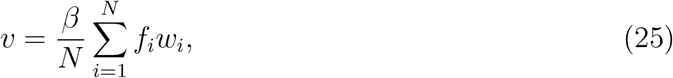

where *β* = *q*|*V_r_*|(*τ_nmda_ + τ_ompo_*). The current per spine *v* is the key dynamical variable in our dimensional reduction procedure and subsequent analysis (see below).

### Dependence of the postsynaptic firing rate *r* on synaptic current *v*

The time scales related to neuronal firing rates and firing adaptation *τ_f_, τ_r_* and *τ_a_* are much faster than the time scale *τ_w_* associated with synaptic plasticity. Therefore, for long times of the order of *τ_w_*, firing rate *r* and postsynaptic current adaptation *s* are in quasi-stationary state, i.e., *dr/dt* ≈ *ds/dt* ≈ 0. This implies a set of coupled algebraic equations:

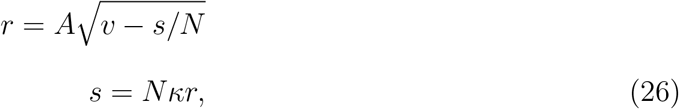

which yields a quadratic equation for *r*, i.e., *r*^2^ + *A*^2^*κr* − *A*^2^*v* = 0. The solution for *r*, which depends on *v*, is given by

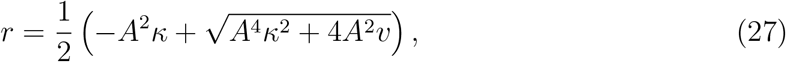

Note that *r* depends nonlinearly on the synaptic current *v*. Additionally, *s/N* is always smaller than *v* in the steady state, which means that *r* in Eq. (26) is well defined.

### Dimensional reduction of the extended BCM model: Dynamic mean-field model

We focus on the population averaged synaptic current *v* (Eq. 25). Since *v* is proportional to weights *w_i_*, and because *r* depends directly on *v*, it is possible to obtain a closed form dynamic equation for plasticity of *v*. Thus, instead of dealing with *N* dimensional dynamics of synaptic weights, we can study a one dimensional dynamics of the population average current *v*. This dimensional reduction is analogous to observing the motion of a center of mass of many particle system, which is easier than simultaneous observation of the motions of all particles. Such an approach is feasible for an analytical treatment where one can directly apply the methods of stochastic dynamical systems and thermodynamics (Van Kampen 2007).

The time derivative of *v*, given by Eq. (25), is denoted with dot and reads

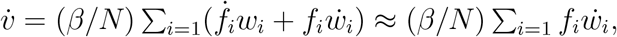

where we used the fact that fluctuations in *f_i_* are much faster than changes in weights *w_i_*, and hence *f_i_* are in stochastic quasi-stationary states. Now, using Eq. (1) for *w_i_* and quasi-stationarity of *θ*, we obtain the following equation for *v*:

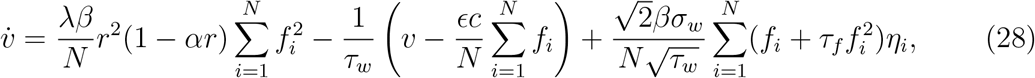

where *c* = *αβ*.

The next step is to perform averaging over fast fluctuations in presynaptic rate *f_i_*. We need to find the following three averages with respect to the random variable

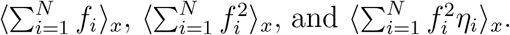

From Eq. (23) it follows that 〈*f_i_*〉_*x*_ = *f_o_*, and thus the first average is

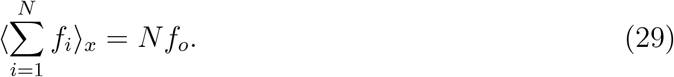

The second average follows from Eq. (24), and we have

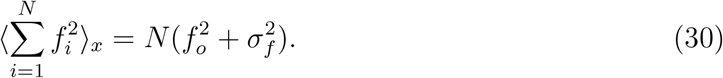

The third average can be decomposed as 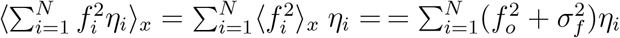, where we used the fact that the noise *η* is independent of the noise *x*, and again Eq. (24).

The final step is to insert the above averages into the equation for 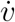 (Eq. 28). As a result we obtain Eq. (3) in the main text, which is a starting point for determining energetics of synaptic plasticity and information characteristics.

### Distribution of synaptic currents in the stochastic mean-field model: weak and strong synapses

Stochastic Eq. (3) for the population averaged synaptic current *v* can be written in short notation as

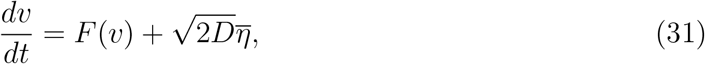

where the function *F*(*v*) is defined as

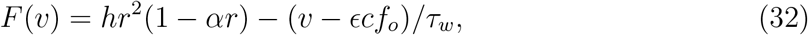

and *D* is the effective noise amplitude (it includes also fluctuations in the presynaptic input) given by

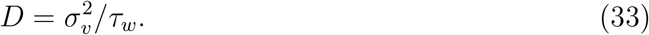

Eq. (31) corresponds to the following Fokker-Planck equation for the probability distribution of the synaptic current *P*(*v|f_o_; t*) conditioned on *f_o_* (Van Kampen 2007):

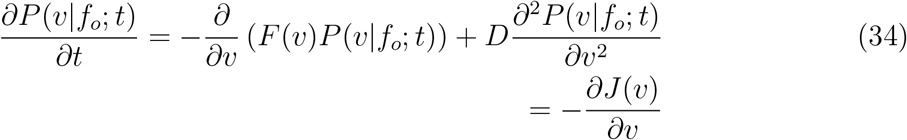

The function *J*(*v*) in the last equality in Eq. (34) is the probability current, which is *J* (*v*) = *F* (*v*) − *D∂P* (*v*)/*∂v*.

The stationary solution of the Fokker-Planck equation (Eq. 34) is obtained for a constant probability current *J* (Gardiner 2004; Van Kampen 2007). For monostable systems, which have a unique steady state (fixed point) one usually sets *J*(*v*) = 0, which corresponds to a detailed balance (Gardiner 2004; Tome 2006). Such a unique steady state corresponds to thermal equilibrium with the environment (Tome 2006) and the solution is of the form (Van Kampen 2007)

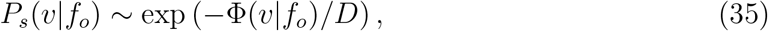

where Φ(*v|f_o_*) is the effective potential for synaptic current *v*, and it is obtained by integration of *F*(*v*) in Eq. (31), i.e.,

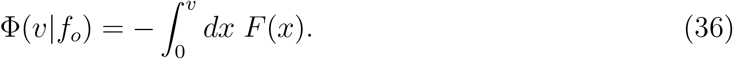

The potential can have either one (monostability) or two (bistability) minima, depending on *f_o_* and other parameters. The explicit form of Φ(*v|f_o_*) is shown in Eq. (13).

For bistable systems, for which there are two possible steady states (fixed points), the situation is more complicated. In the presence of nonzero, *D* > 0, effective synaptic noise, there can be noise induced jumps between the two fixed points. In this case the probability current J in the steady state must be a nonzero constant, because of the exchange of probabilities between the two fixed points, or equivalently, because of the stochastic jumps of *v* between two potential wells (Gardiner 2004). For small noise *D*, such jumps between the two potential wells happen on very long time scales. This longtime dynamics is the primary reason that the stationary state of such “driven” systems (by thermal and presynaptic fluctuations) is globally out of thermal equilibrium with the environment and it is called thermodynamic nonequilibrium steady state, in which the detailed balance is broken (Tome 2006). However, locally, close to each fixed point and for not too long times the system is in local thermal equilibrium. Thus, we can locally approximate the probability distribution of *v* by the form given in Eq. (35), by expanding the potential Φ_*s*_(*v*) around *v_d_* and *v_u_*. Using a Gaussian approximation, which should be valid for small *D* (either for large *N* or for small *σ_w_* or both), we can write *P_s_*(*v*) for *v* close to *v_d_* as 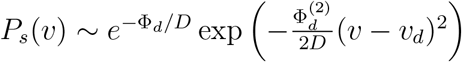, and for *v* close to *v_u_* we have 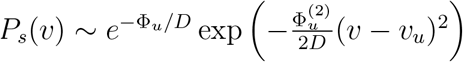, where 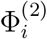 is the second derivative of the potential with respect to *v* at *v_i_*, where the subscript *i* is either *d* (down state) or *u* (up state). For the sake of computations we have to extend these local approximations to longer intervals of *v*, corresponding to the domains of attraction for two fixed points *v_d_* and *v_u_*. Consequently, we assume that the first approximation works for 0 ≤ *v* ≤ *v_max_*, and the second for *v* > *v_max_*. In sum, we approximate the stationary probability density *P_s_*(*v|f_o_*) as two Gaussian peaks centered at *v_d_* and *v_u_*:

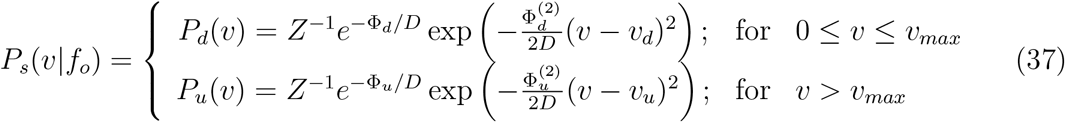

where *Z* is the normalization factor, which can be written as a sum *Z = Z_d_ + Z_u_*, with

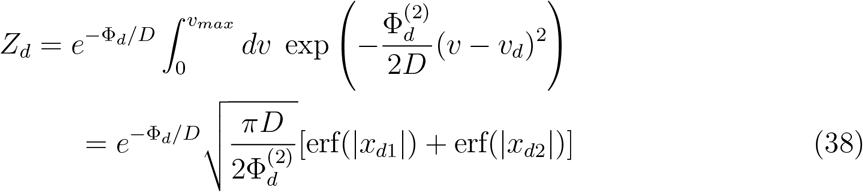

and

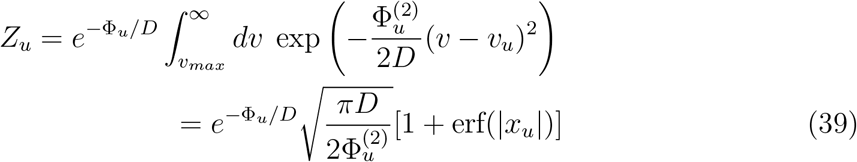

where erf(…) is the error function, 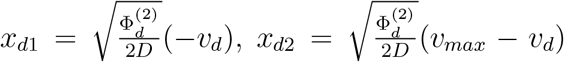, and 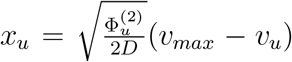. Note that because the unstable fixed point *v_max_* depends on *f_o_*, the arguments of the error functions in and change with changes in *f_o_*. This influences the determination of *p_d_,p_u_*, as well as energy rate and Fisher information (see below).

### Fractions of weak and strong synapses

We define the fraction of synapses in the down state (fraction of weak synapses) as the probability that synaptic current *v* is in the domain of attraction of the down fixed point in the deterministic limit. This takes place for *v* in the range 0 ≤ *v* ≤ *v_max_*, where *v_max_* is the unstable fixed point separating the two stable fixed points *v_d_* and *v_u_*. By analogy, the fraction of synapses in the up state is the probability that *v* is greater than *v_max_*. We can write this mathematically as

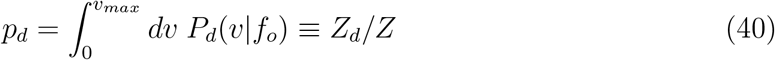

and

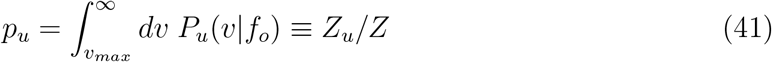

where *P_d_*(*v|f_o_*) and *P_u_*(*v|f_o_*) are given by Eq. (37). Using the expressions for and *Z_u_*, we find an explicit form of as

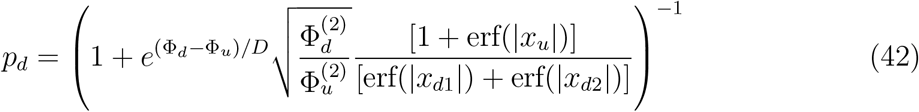

Note that *p_d_* and *p_u_* sum to unity, since *Z = Z_d_ + Z_u_*.

### Average values of *v* and *r* in the mean-field

Average value of the synaptic current *v* in the mean-field is denoted as 〈*v*〉, and computed as

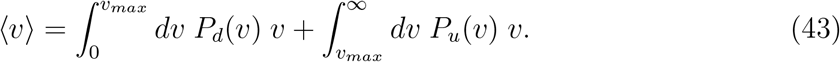

Execution of these integrals yields

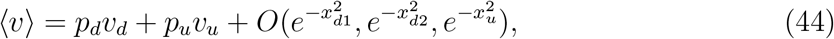

where 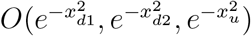 denotes small exponential terms in the limit of very small *D*.

Standard deviation of *v* can be found analogically, which yields

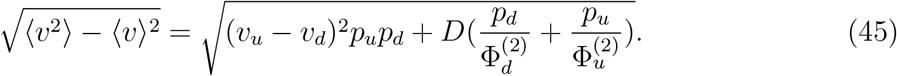

The average value of the postsynaptic firing rate *r*, denoted as 〈*r*〉, is computed in the limit of large *A* (see Eq. 6). In this limit *r* ≈ (*v/κ*) [1 − *v*/(*Aκ*)^2^], and we find

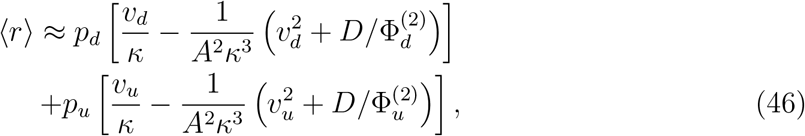

which means that the form of 〈*r*〉 is more complicated than 〈*v*〉.

### Transitions between weak and strong synaptic states: Kramer escape rate

For cortical neurons the number of spines per neuron are very large, i.e. *N* ~ 10^3^ − 10^4^ (Elston et al 2001; DeFelipe et al 2002; Sherwood et al 2020), and thus one can expect that *σ_v_* is small and consequently the fluctuations around the population average current *v* are rather weak. The results described below are obtained in the limit of small *σ_v_*.

Plastic synapses can jump between down and up states due to effective synaptic noise *σ_v_* or *D*. From a physical point of view, this corresponds to a noise induced “escape” of some synapses through a potential barrier. Average dwelling times in the up (*T_u_*) and down (*T_d_*) states can be determined from the Kramers’s formula (Van Kampen 2007):

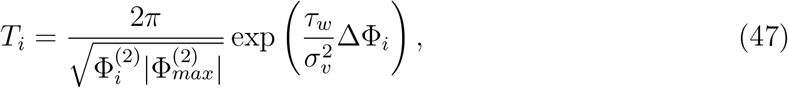

where the index *i* = *d* or 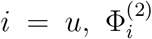 and 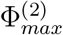 are the second derivatives of the potential at its minima (*v* = *v_i_*) and maximum (*v* = *v_max_*), and the potential difference ΔΦ¿ = Φ(*v_max_*) − Φ(*v_i_*) > 0. Note that for large number of synapses *N*, the exponential factor in Eqs. (47) can be large, which can lead to very long dwelling times that are generally much longer than any time scale in the original Eqs. (1–2). The fact that the times *T_u_* and *T_d_* are long but finite is an indication of metastability of “locally” stable up and down synaptic states.

There exist a relationship between fractions of weak/strong synapses and the Kramer’s escape times *T_d_* and *T_u_* in the limit of very weak noise *D* ↦ 0. Namely, it can be easily verified that in this limit 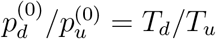, and consequently, we can write

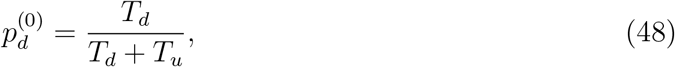

where 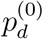 is the fraction of weak synapses for *D* ↦ 0 given by

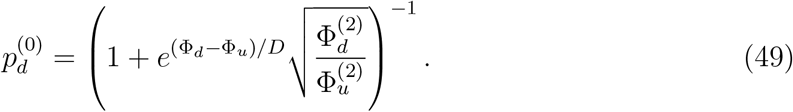

### Memory lifetime

Synaptic memory lifetime *T_m_* is defined as a characteristic time the synapses remember a perturbation to their steady state distribution. Mathematically, it means that we have to consider a time-dependent solution of the probability density *P*(*v|f*_0_; *t*) to the Fokker-Planck equation given by Eq. (34). This solution can be written as (van Kampen 2007; Risken 1996)

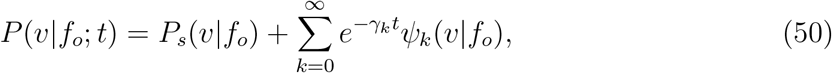

where *γ_k_* and *ψk*(*v|f_o_*) are appropriate eigenvalues and eigenvectors. The eigenvalues are inverses of characteristic time scales, which describe a relaxation process to the steady state. The smallest eigenvalue, denoted as *γ*_0_, determines the longest relaxation time 1/*γ*_0_, and we associate that time with the memory lifetime It has been shown that *γ*_0_ = 1/*T_d_* + 1/*T_u_* (van Kampen 2007; Risken 1996), which implies that

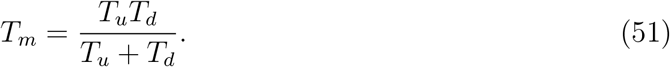

A similar approach, through eigenvalues, to estimating the memory lifetime was adopted also in (Fusi and Abbott 2007).

### Entropy production rate, entropy flux, and power dissipated by plasticity

Processes underlying synaptic plasticity (e.g. AMPA receptor trafficking, PSD protein phosphorylation, as well as protein synthesis and degradation; see Huganir and Nicoll (2013), Choquet and Triller (2013)) operate out of thermodynamic equilibrium, and therefore require energy influx. At a stochastic steady state, this energy is dissipated as heat, which roughly corresponds to a metabolic rate of synaptic plasticity. The rate of dissipated energy is proportional to the average rate of decrease in the effective potential Φ, or equivalently to the entropy production rate (Nicolis and Prigogine 1977).

Given the above, we can write the energy rate for synaptic plasticity *Ė* as 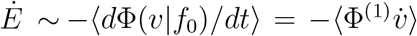, where Φ^(1)^ is the first derivative of Φ with respect to *v*, the symbol 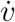 is the temporal derivative of v, and the averaging 〈…〉 is performed over the distribution *P*(*v|f*_0_). The second equality follows from the fact that *v* is the only variable in the potential that changes with time on the time scale *τ_w_*. Next, we can use Eq. (3) or (31) in the equivalent form, namely 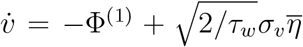, and this equation resembles the motion of an overdamped particle (with negligible mass) in the potential Φ, with *v* playing the role of a spatial coordinate. After that step, we can write the energy rate as 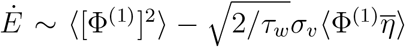. The final step is to use the Novikov theorem (Novikov 1965) for the second average, i.e. 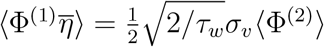. This leads to

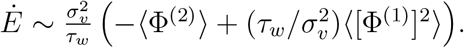

We can obtain a similar result for *Ė* using a thermodynamic reasoning. The dynamics of synaptic plasticity is characterized by the distribution of synaptic currents per synapse *P*(*v|f_o_*), which evolves in time according to Eq. (34). With this distribution we can associate the entropy *S*(*t*), defined as 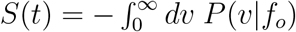 ln *P*(*v|f_o_*), measuring the level of order in a typical spine. It can be shown (Nicolis and Prigogine 1977; Tome 2006; Tome and de Oliveira 2010) that the temporal derivative of the entropy, dS/dt, is composed of two competing terms, *dS/dt* = Π − Γ, called entropy production rate (Π) and entropy flux (Γ), both per synapse. In the case of thermodynamic equilibrium, which is not biologically realistic, one has *dS/dt* = Π = Γ = 0, and there is neither energy influx to a system nor dissipated energy to the environment. However, for processes out of thermodynamic equilibrium, relevant for spine dynamics, we still can find a stationary regime where entropy of the spine does not change, *dS/dt* = 0, but entropy flux Γ and entropy production Π are nonzero and balance each other (Nicolis and Prigogine 1977; Tome 2006). It is more convenient to determine the stationary dissipated power by finding the entropy flux, which is given by (Tome 2006; Tome and de Oliveira 2010) (see Suppl. Infor.)

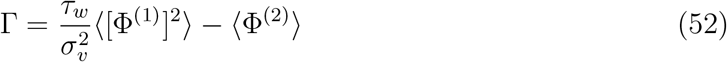

Note that Eq. (52) is very similar in form to the energy rate *Ė* derived above; the two expressions differ only by the factor 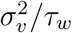, and none of them has the units of energy (Γ has the unit of the inverse of time). Thus, we need to introduce the energy scale in the problem. Generally, the stationary dissipated power per synapse *Ė* can be written as *Ė* = *Ė_o_*Π = *Ė_o_*Γ (Nicolis and Prigogine 1977), where *Ė_o_* is the characteristic energy scale associated with spine conductance changes, and its value is estimated next.

### Estimation of the characteristic energy scale for synaptic plasticity

As was said in the Introduction, the BCM model (either classical or extended) is only a phenomenological model of plasticity that does not relate directly to the underlying molecular processes in synapses. Consequently, a small, single, change of synaptic weight by Δ*w_i_* in Eq. (1) is accompanied in reality by many molecular transitions in synapse i. This means that a single degree of freedom related to *w_i_* is in fact associated with many, hidden, molecular degrees of freedom. To be realistic in our energy cost estimates, we have to include those hidden degrees of freedom.

If we dealt with a process representing a single degree of freedom, then the energy scale *E_o_* relating entropy flux Γ and energy rate *Ė*, would be *E_o_* = *kT* (Nicolis and Prigogine 1977), where *k* is the Boltzmann constant and *T* is the tissue absolute temperature (*T* ≈ 310 K). However, a dendritic spine is a composite object with multiple components and many degrees of freedom (Bonhoeffer and Yuste 2002; Holtmaat et al 2005; Meyer et al 2014; Choquet and Triller 2013), and hence the characteristic energy scale *E_o_* is much bigger than kT. The changes in spine conductance on time scale of ~ 1 hr, i.e. for e-LTP and e-LTD, are induced by protein interactions in PSD (Lisman et al 2012; Kandel et al 2014) and subsequent membrane trafficking associated with AMPA and NMDA receptors (Borgdorff and Choquet 2002; Huganir and Nicoll 2013; Choquet and Triller 2013). Protein interactions are powered by phosphorylation process, which is one of the main biochemical mechanism of molecular signal transduction in PSD rele-vant for synaptic plasticity (Bhalla and Iyengar 1999; Zhu et al 2016). Phosphorylation rates in an active LTP phase can be very fast, e.g., for CaMKII autophosphorylation they are in the range 60 − 600 min^−1^ (Bradshaw et al 2002). Other processes in a spine, most notably protein turnovers in PSD (likely involved in l-LTP and l-LTD), are much slower ~ 3.7 days (Cohen et al 2013), and therefore their contribution to the energetics of the early phase of spine plasticity seems to be much less important (see, however Discussion for an estimate of the protein turnover energy rate).

The energy scale for protein interaction can be estimated as follows. A typical dendritic spine contains about 10^4^ proteins (including their copies) (Sheng and Hoogenraad 2007). One cycle of protein phosphorylation requires the hydrolysis of 1 ATP molecule (Hill 1989; Qian 2007), which costs about 20*kT* (Phillips et al 2012). Each protein has on average 4-6 phosphorylation sites (Collins et al 2005; Trinidad et al 2012). If we assume conservatively that only about 20% of all PSD proteins are phosphorylated, then we obtain the energy scale for protein interactions roughly 2 · 10^5^*kT*, which is 8.6 · 10^−16^ J.

Energy scale for receptor trafficking can be broadly decomposed into two parts: energy required for insertion of the receptors into the spine membrane, and energy related to their horizontal movement along the membrane to the top near a presynaptic terminal. The insertion energy for a typical protein is either about 3 − 17 kcal/mol (Gumbart et al 2011) or 8 − 17*kT* (Grafmuller et al 2009), with the range spanning 4 − 25*kT*, and is caused by a deformation in the membrane structure (Gumbart et al 2011). Since an average spine contains about 100 AMPA (Matsuzaki et al 2001;Smith et al 2003) and 10 NMDA (Nimchinsky et al 2004) receptors, we obtain the total insertion energy in the rage 500 − 3200*kT*. The second, movement contribution can be estimated by noting that typical forces that overcome friction and push macromolecules along membrane are about 10 pN, and they are powered by ATP hydrolysis (Fisher and Kolomeisky 1999). AMPA and NMDA receptors have to travel a spine distance of about 1 *μ*m (Benavides-Piccione et al 2013), which requires the work of 110 · 10^−11^ · 10^−6^ N·m= 1.1 · 10^−15^ J or 2.5 · 10^5^kT. The latter figure is 100 times larger than the insertion contribution, which indicates that the energy scale for receptor trafficking is dominated by the horizontal movement and is similar to the above for protein phosphorylation.

To summarize, the total energy scale *E_o_* for spine conductance is about *E_o_* = 2 · 10^−15^ J, or equivalently 4.6 · 10^5^*kT* (or 2.3 · 10^4^ ATP molecules).

### Analytical approximation of the energy rate related to synaptic plasticity

It is not possible to find analytically the entropy flux Γ in Eq. (52) for an arbitrary probability distribution. However, Γ can be determined approximately for the probability distribution *P_s_*(*v|f_o_*) in Eq. (37), by the saddle point method as a series expansion in the small noise amplitude *D*, which is proportional to 1/*N*. We can write the entropy flux Γ in terms of the probability densities *P_d_* and *P_u_* appearing in Eq. (37) as

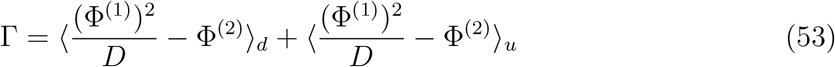

where

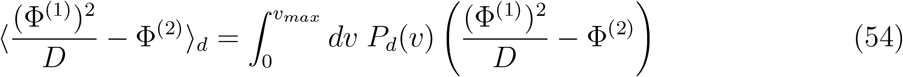

and

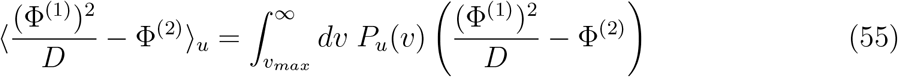

The essence of the saddle point method is in noting that for very small *D*, the probability distributions in Eq. (37) have two sharp maxima corresponding to two most likely synaptic currents *v_d_* and *v_u_*. This implies that the values of *v* that are the closest to *v_d_* and *v_u_* in (Φ^(1)^)^2^/*D* − Φ^(2)^ provide the biggest contributions to the integrals in Eqs. (54) and (55), and hence to the entropy flux Γ. Consequently, we have to expand the function (Φ^(1)^(*v*))^2^/*D* − Φ^(2)^ (*v*) around *v_d_* and *v_u_*.

For *v* near *v_d_*, the expansion is simpler if we introduce a unitless variable *x*, related to *v* such that 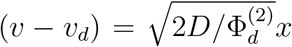, where 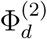 is the second derivative of Φ(*v*) at *v* = *v_d_*. Then to the order ~ *D* we have:

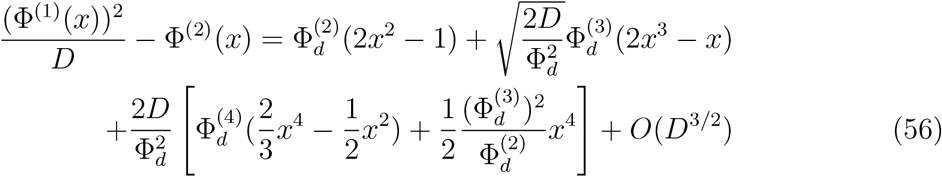

A similar expression holds *v* near *v_u_*, with a substitution 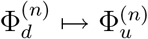. Thus for 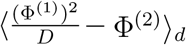 we have

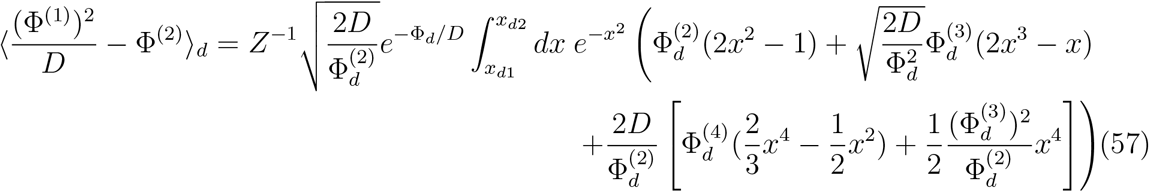

where the limits of integration are 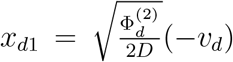, and 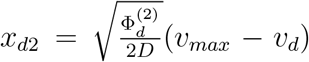. Execution the above integrals yields

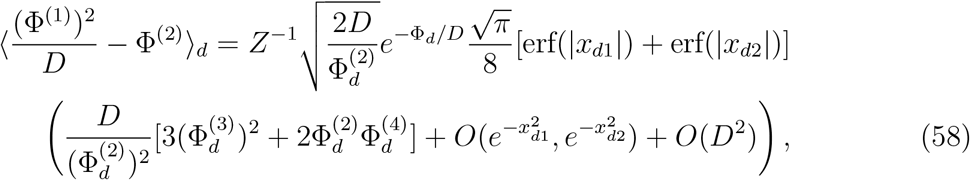

where in the limit of small *D*, the exponential terms 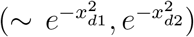 are small, and thus negligible. Next, it is easy to note that the prefactor in front of the large bracket simplifies, i.e, 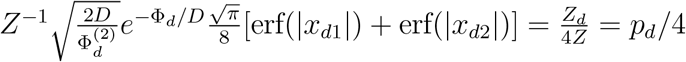.

Applying the same procedure for 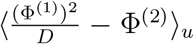 gives us the total expression for the entropy flux Γ

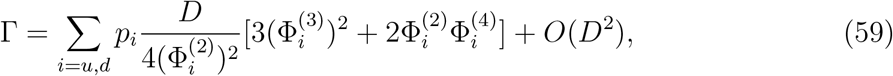

where *i* = *d* (down state) or *i = u* (up state).

Having the entropy flux, we can determine analytically the power dissipated per synapse *Ė* due to synaptic plasticity. The result is

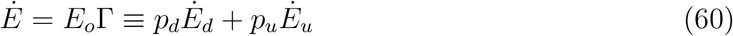

where *Ė_d_* and *Ė_u_* are the energy rates dissipated in the down and up states, respectively. They take the form:

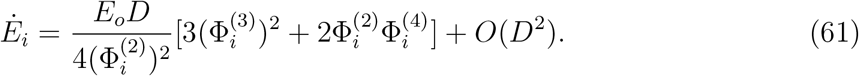

Note that the first nonzero contribution to the energy rate is of the order ~ *D*.

### Neuron energy rate related to fast electric signaling

We provide below an estimate of the energy used by a sensory neuron for short-term signaling for the sake of comparison with the energy requirement of synaptic plasticity. It has been suggested that the majority of neuronal energy goes to pumping out Na^+^ ions (Na^+^-K^+^-ATPase), which accumulates mostly due to neural spiking activity, synaptic background activity, and passive Na^+^ influx through sodium channels at rest (Attwell and Laughlin 2001). It has been shown that this short-term neuronal energy cost can be derived from a biophysical neuronal model, compared across species, and represented by a relatively simple equation (Karbowski 2009, 2012):

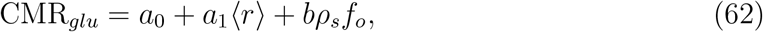

where CMR_*glu*_ is the glucose metabolic rate [in *μ*mol/(cm^3^· min)], *ρ_s_* is the synaptic density, 〈*r*〉 is the average postsynaptic firing rate, and the parameters *a*_0_, *a*_1_, and *b* characterize the magnitude of the above three contributions to the neural metabolism, i.e. resting, firing rate, and synaptic transmission, respectively (Karbowski 2012). The average postsynaptic rate (r) is found from Eq. (46).

According to biochemical estimates, one oxidized glucose molecule generates about 31 ATP molecules (Rolfe and Brown 1997). In addition, 1 ATP molecule provides about 20kT of energy (Phillips et al 2012). This means that the short-term energy rate per neuron, denoted as *Ė_n_*, is given by

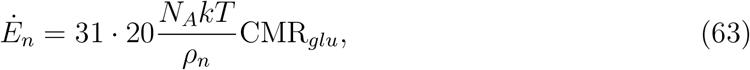

where *N_A_* is the Avogadro number, and *ρ_n_* is the neuron density. We estimate the ratio of the synaptic plasticity power to neural power, i.e. *Ė/Ė_n_* across different presynaptic firing rates for three areas of the adult human cerebral cortex (frontal, temporal, and visual), and two areas of macaque monkey cerebral cortex (frontal and visual).

The values of the parameters *a*_0_ and *α*_1_ in Eq. (62) are species- and area-independent, and they read *a*_0_ = 2.1 · 10^−1^° mol/(cm^3^ s), and *a*_1_ = 2.3 · 10^−9^ mol/cm^3^ (Karbowski 2012). The rest of the parameters take different values for human and macaque cortex. Most of them are taken from empirical studies, and are given below. The parameter b, present in Eq. (62), is proportional to the neurotransmitter release probability and synaptic conductance, and it was estimated based on fitting developmental data for glucose metabolism CMR_*glu*_ and synaptic density *ρ_s_* (which vary during the development) to the formula (62) (Karbowski 2012).

The following data are for an adult human cortex. The adult CMR_*glu*_ is 0.27 μmol/(cm^3^·min) (frontal cortex), 0.27μmol/(cm^3^·min) (visual cortex), and 0.24 μmol/(cm^3^·min) (temporal cortex) (Chugani 1998). The parameter b reads: 1.16 · 10^−20^ mol (frontal), 0.63 · 10^−20^ mol (visual), 0.17 · 10^−20^ mol (temporal) (Karbowski 2012). Note that the value of b is 7 times larger for the frontal cortex than for the temporal, which might suggest that the product of neurotransmitter release probability and synaptic conductance is also 7 fold larger in the frontal cortex. This high difference may seem unlikely, however, it is still plausible, given that the release probability is highly variable and can assume values between 0.05-0.7 (Bolshakov and Siegelbaum 1995, Frick et al 2007; Volgushev et al 2004; Murthy et al 2001), and synaptic weights in the cortex are widely distributed (Loewenstein et al 2011). Neuron density *ρ_n_* reads: 36.7· 10^6^ cm^−3^ (frontal), 66.9 · 10^6^ cm^−3^ (visual), 59.8 · 10^6^ cm^−3^ (temporal) (Pakkenberg and Gundersen 1997). Synaptic density *ρ_s_* reads: 3.4 · 10^11^ cm^−3^ (frontal), 3.1 · 10^11^ cm^−3^ (visual), 2.9 · 10^11^ cm^−3^ (temporal) (Huttenlocher and Dabholkar 1997).

The following data are for an adult (6 years old) macaque monkey cortex. The adult CMR_*glu*_ is 0.34 μmol/(cm^3^·min) (frontal cortex), 0.40 μmol/(cm^3^·min) (visual cortex)(Noda et al 2002). The parameter b reads: 0.4 · 10^−2^° mol (frontal), and 3.8 · 10^−2^° mol (visual) (Karbowski 2012). Neuron density *ρ_n_* reads: 9 · 10^7^ cm^−3^ (frontal), 31.9 · 10^7^ cm^−3^ (visual) (Christensen et al 2007). Synaptic density reads: 5· 10^11^ cm^−3^ (frontal) (Bourgeois et al 1994), 6 · 10^11^ cm^−3^ (visual) (Bourgeois and Rakic 1993).

### Fisher information and coding accuracy in synapses

Fisher information *I_F_*(*f_o_*) about the driving input *f_o_* is a good approximation of the mutual information between the driving presynaptic activity and postsynaptic current *v* (Brunel and Nadal 1998). It is also a measure of the coding accuracy and it is defined as (Cover and Thomas 2006)

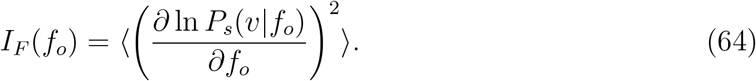

Taking into account the form of probability density, Eq. (37), we can rewrite this equation as

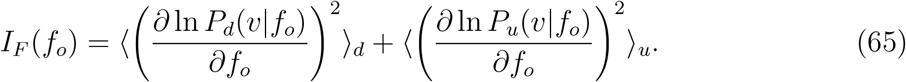

where

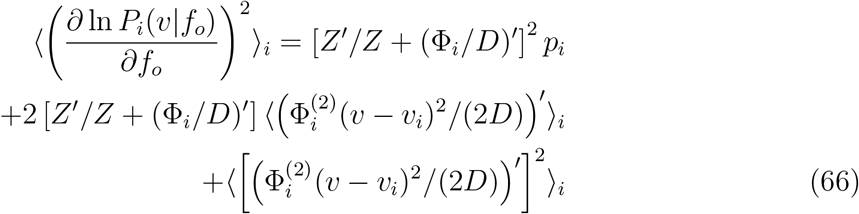

Our first goal is to express the factor *Z’/Z* in terms of the potential and its derivatives.

To do it, we compute the following average:

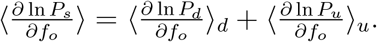

The left hand side of this equation is zero, since

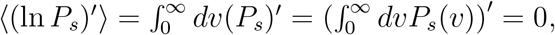

where a prime denotes a derivative with respect to *f_o_*. Additionally,

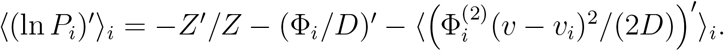

Combining the last two equations we obtain a relation between *Z’/Z* and the potentials:

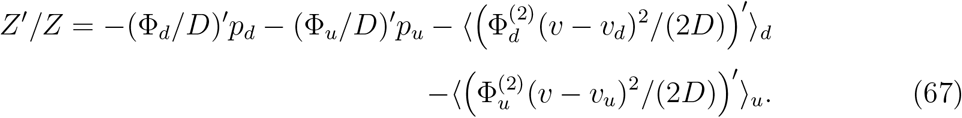

After insertion of this expression into Eq. (66) and after some algebra, we arrive at the Fisher information

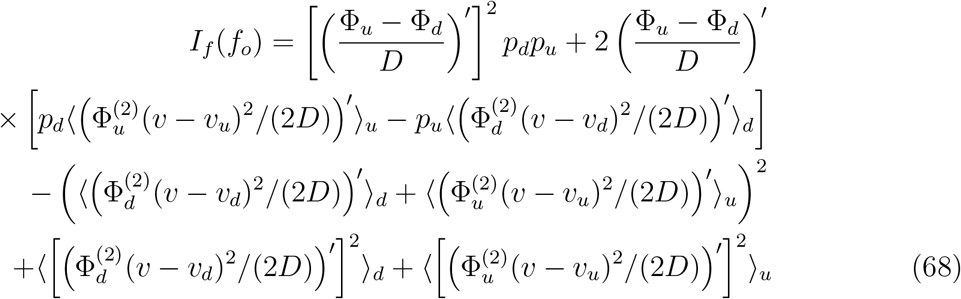

The averages in the above equation can be computed to yield:

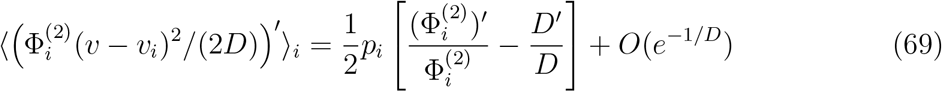

and

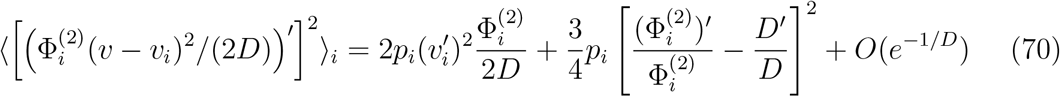

After insertion of these expressions into Eq. (68), and some algebraic manipulations, we arrive at Eq. (17) for in the Results.

### Relationship between synaptic energy rate and Fisher information in the limit *D* ↦ 0

Below we derive the relation given by Eqs. (18) and (19) in the limit of very weak synaptic noise, *D* ↦ 0. In this limit, it can be noted that the product of the fractions of weak and strong synapses is

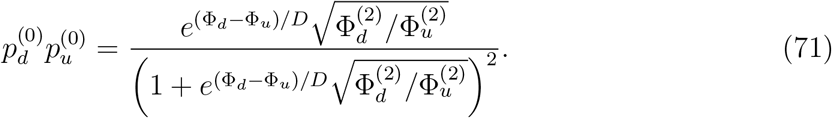

This expression enables us to write in a compact form the derivative of 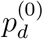 with respect to *f_o_* as

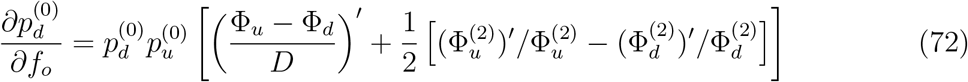

where the prime denotes differentiation with respect to *f_o_*. On the other hand it can be noted that, in the bistable regime, the Fisher information in Eq. (17) in the leading order 1/*D*^2^ can be written as

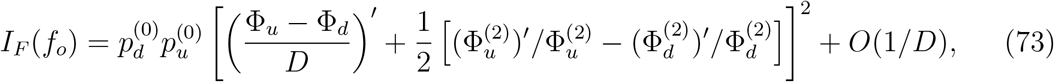

which is similar in form to the expression for 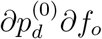. This suggests that we can combine the two equations, and arrive at

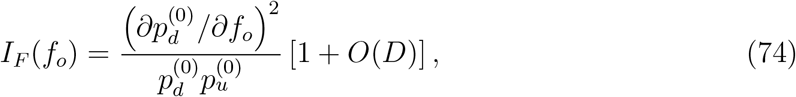

which is Eq. (18) in the Results.

Next, we want to relate Eq. (74) for *I_F_* to the energy rate. Energy rate 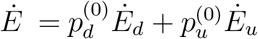 can be differentiated with respect to *f_o_*, which yields

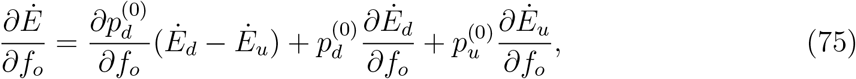

where we used the relation 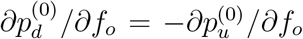, which follows from the fact that 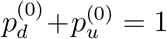. Now it is crucial to note that the first term in 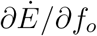, i.e. 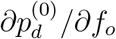 is of the order 1/*D*, whereas the rest terms 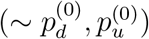 are of the order of one. This implies that the first term dominates in the limit *D* ↦ 0. Thus, we can write approximately the expression for 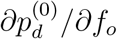, involving the energy rate as

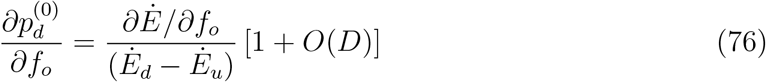

Finally, if we combine Eqs. (74) and (76), we obtain Eq. (19) in the Results for the bistable regime.

### Numerical simulations of the full synaptic system

Numerical stochastic dynamics of the whole synaptic system given by Eqs. (1–2) were performed using a stochastic version of Runge-Kutta scheme (Roberts 2012).

Energy dissipated for plasticity by the full synaptic system (Eqs. 1–2) was computed numerically using the approach presented in (Tome 2006, Tome 2010). We can rewrite Eq. (1) in a more compact form as

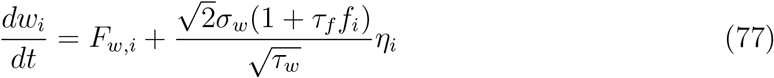

where

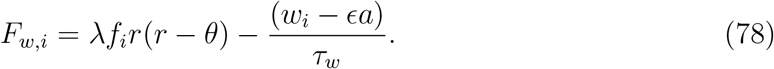

This enables us to write the entropy flux in the steady state (equivalent to entropy production rate) of the full synaptic system in a compact form. Consequently, the numerical entropy flux per synapse of the whole system Γ_*num*_ is

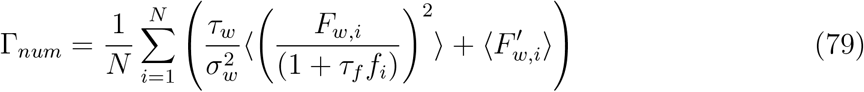

where 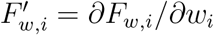, and it is given by

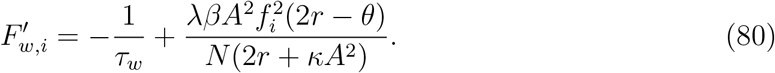

The numerical energy rate *Ė_num_* is *Ė_num_* = *E_o_*Γ_*num*_. The brackets 〈…〉 in Eq. (79) denote averaging over fluctuations in synaptic noise and presynaptic firing rates (averaging over *η* and *x* stochastic variables). In numerical simulations, these averages are computed as temporal averages over long simulation time. This equivalence in averaging is guaranteed due to ergodic theorem. The minimal number of time steps for numerical convergence is of the order of ~ 10^5^.

### Parameters used in computations

The following values of various parameters were used: *V_r_* = −65 mV, *q* = 0.35 (Vol-gushev et al 2004), *τ_nmda_* = 150 msec (Nimchinsky et al 2004), *τ_ompo_* = 5 msec (Smith et al 2003), *τ_f_* = 1.0 sec, *a* = 1.0 nS, *α* = 0.3 sec (Zenke et al 2013), *ϵ* = 3 · 10^−4^, 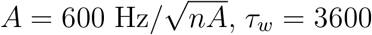 sec (Frey and Morris 1997; Zenke et al 2013), *σ_f_* = 10 Hz (Buzsaki and Mizuseki 2014), *N* =2 · 10^3^ (average value for many species of primates; see Sherwood et al (2020), Elston et al (2001)). The amplitude of synaptic weight noise *σ_w_* was taken in the range 0.02 ≤ *σ_w_* ≤ 0.5 nS, which is the range suggested in experimental studies (Matsuzaki et al 2001; Smith et al 2003). The two undetermined parameters are λ and *κ*, and two sets of values were used for them: (i) *κ* = 0.001 (nA·sec), λ = 9 · 10^−7^ (nS·sec^2^), and (ii) *κ* = 0.012 (nA·sec), λ = 10^−5^ (nS·sec^2^), inorder to obtain a transition to the bistable regime for *f_o_* ~ 1 − 5 Hz. The value of *A* was chosen to have postsynaptic firing rate in the range 0.1 − 10 Hz. The value of *κ* was chosen to obtain *v_u_* in the neurophysiological range ~ 1 pA (O’Connor et al 2006).

## Supporting Information

**S1 Text.** This file contains the details of some calculations. (PDF)

## Acknowledgments

The work was supported by the Polish National Science Centre (NCN) grant no. 2015/17/B/NZ4/02600.

