## Supplemental Information for "Energetics of stochastic BCM type synaptic plasticity and storing of accurate information"

Jan Karbowski

*Institute of Applied Mathematics and Mechanics,*

*University of Warsaw, ul. Banacha 2, 02-097 Warsaw, Poland;*

#### 1 Determination of the fixed points of Eq. (3).

Fixed points of Eq. (3) in the main text in the limit $N \mapsto \infty$, i.e. without noise, are determined from the condition $dv/dt = 0$. Since, the variables $v$ and $r$ are monotonically related by Eq. (6) in the main text, we can invert this formula and write equivalently:

$$v = \kappa r + \left(\frac{r}{A}\right)^2. \tag{1}$$

This step enables us to solve equation $dv/dt = 0$ not in $v$ variable but in $r$ variable, which is much easier. Thus, the fixed points of Eq. (3) in the main text are determined from the following equation:

$$h\tau_w r^2(1-\alpha r) - \left(\kappa r + \frac{r^2}{A^2} - \epsilon c f_o\right) = 0. \tag{2}$$

We can rearrange this equation and obtain a polynomial of the third degree in $r$:

$$\alpha h\tau_w r^3 - \left(h\tau_w - \frac{1}{A^2}\right) r^2 + \kappa r - \epsilon c f_o = 0. \tag{3}$$

We can solve this equation iteratively, using an $\epsilon$ expansion, i.e., we look for the solution in the form: $r = r_0 + r_1\epsilon + r_2\epsilon^2 + O(\epsilon^3)$, where $\epsilon \ll 1$, and $r_0, r_1, r_2$ are unknowns to be determined. This solution corresponds to the case when basal conductance of the synapse/spine is very small (thin spines are weakly conducting electric current). After the above substitution for $r$, Eq. (3) becomes:

$$R_0 + R_1\epsilon + R_2\epsilon^2 + O(\epsilon^3) = 0, \tag{4}$$

where "the coefficients" $R_0, R_1, R_2$ near different powers of $\epsilon$ given below must vanish, i.e.,

$$R_0 = -\alpha h\tau_w r_0^3 + \left(h\tau_w - \frac{1}{A^2}\right) r_0^2 - \kappa r_0 = 0. \tag{5}$$

$$R_1 = -3\alpha h\tau_w r_0^2 r_1 + 2\left(h\tau_w - \frac{1}{A^2}\right) r_0 r_1 - \kappa r_1 + cf_o = 0. \tag{6}$$

$$R_2 = -3\alpha h\tau_w r_0(r_0 r_2 + r_1^2) + \left(h\tau_w - \frac{1}{A^2}\right)(r_1^2 + 2r_0 r_2) - \kappa r_2 = 0. \tag{7}$$

Solution of the system (5-7) in terms of $r_0, r_1, r_2$ leads to three different solutions in terms of $r$ (three states: down (d), up (u), and intermediate (max)) which read:

$$r_d = \frac{cf_o}{\kappa}\epsilon + \frac{(cf_o)^2(h\tau_w - 1/A^2)}{\kappa^3}\epsilon^2 + O(\epsilon^3), \tag{8}$$

$$r_u = r_+ + \frac{cf_o}{[3\alpha h\tau_w r_+^2 - 2(h\tau_w - 1/A^2)r_+ + \kappa]}\epsilon + O(\epsilon^2) \tag{9}$$

and

$$r_{max} = r_- + \frac{cf_o}{[3\alpha h\tau_w r_-^2 - 2(h\tau_w - 1/A^2)r_- + \kappa]}\epsilon + O(\epsilon^2) \tag{10}$$

where $r_+$ and $r_-$ are given by

$$r_\pm = \frac{(h\tau_w - 1/A^2) \pm \sqrt{\Delta}}{2\alpha h\tau_w}, \tag{11}$$

with $\Delta = (h\tau_w - A^{-2})^2 - 4\alpha\kappa h\tau_w$. The condition for the existence of $r_u$ and $r_{max}$ is $\Delta \geq 0$. If we use Eq. (1) again, we obtain $v_d, v_u, v_{max}$ in the leading order in $\epsilon$, which are present in the main text.

#### 2 Entropy production and entropy flux for nonequilibrium systems.

In this section we derive the expressions for entropy production and entropy flux, which is given by Eq. (52) in the main text.

Eq. (34) in the main text can be written as:

$$\frac{\partial P(v|f_o)}{\partial t} = -\frac{\partial J}{\partial v}, \tag{12}$$

where $J$ is the probability flux given by

$$J(v) = F(v)P(v|f_o) - \frac{\sigma_v^2}{\tau_w}\frac{\partial P(v|f_o)}{\partial v}. \tag{13}$$

The temporal derivative of the entropy $S$, defined as $S = -\int dv P(v|f_o) \ln P(v|f_o)$, becomes after using Eq. (12) above as follows [1]:

$$\frac{\partial S}{\partial t} = \int dv [1 + \ln P(v|f_o)] \frac{\partial J}{\partial v}. \tag{14}$$

The integral can be done by parts (using the boundary condition that $P$ and its derivative are 0 at the boundaries), and hence

$$\frac{\partial S}{\partial t} = -\int dv J(v) \frac{1}{P(v|f_o)} \frac{\partial P}{\partial v}. \tag{15}$$

Form Eq. (13) above, we get

$\frac{1}{P}\frac{\partial P}{\partial v} = \frac{\tau_w}{\sigma_v^2}\Big(F - \frac{J}{P}\Big)$,

and after substituting this expression for $\partial P/(P\partial v)$ in Eq. (15), and performing rearrangements, we obtain $dS/dt = \Pi - \Gamma$, where the entropy production $\Pi$ and entropy flux $\Gamma$ are given by

$$\Pi = \frac{\tau_w}{\sigma_v^2}\int dv \frac{J(v)^2}{P(v|f_o)}, \tag{16}$$

and

$$\Gamma = \frac{\tau_w}{\sigma_v^2}\int dv J(v)F(v). \tag{17}$$

Entropy production is a measure of how fast energy is dissipated in the system (in units $sec^{-1}$) in the form of heat. At a nonequilibrium steady state (considered in the paper), both $\Pi$ and $\Gamma$ are nonzero and equal to each other. Thus, to have entropy production rate at the steady state, it is enough to determine $\Gamma$, since $\Gamma$ is easier to compute.

Let us find a more useful form of the entropy flux. For this we insert Eq. (13) for $J$ into Eq. (17), and decompose the resulting integral into two integrals, one with $P$ and second with $\partial P/\partial v$. The second integral is done by parts, and we obtain the entropy flux equivalently as:

$$\Gamma = \frac{\tau_w}{\sigma_v^2}\int dv P(v|f_o)F(v)^2 + \int dv P(v|f_o)\frac{\partial F}{\partial v}. \tag{18}$$

Since the function $F(v)$ is directly related to the derivative of the potential $\Phi(v|f_o)$ with respect to $v$, i.e. $F(v) = -\Phi^{(1)}(v|f_o)$, we reproduce Eq. (52) in the main text.

### 3 Saddle point approximation.

In this section it is shown how the saddle point approximation works. Let's consider the following integral with an arbitrary smooth function $G(v)$:

$$I = \int_0^\infty dv G(v) e^{-(v-v_o)^2/2\sigma^2}. \tag{19}$$

The idea is that for small $\sigma$ the biggest contribution to the integral comes from $v$ that are close to $v_o$. Specifically, we change a variable of integration from $v$ to a new variable $x = (v - v_o)/\sqrt{2\sigma^2}$. Thus, our integral $I$ in the new variable $x$ takes the form

$$I = \sqrt{2\sigma^2} \int_{x_0}^\infty dx G(v_o + \sqrt{2\sigma^2} x) e^{-x^2}, \tag{20}$$

where the lower limit of integration is $x_0 = -v_o/\sqrt{2\sigma^2} < 0$. The next step is to expand the function $G$ in powers of $x$ and to compute the emerging integrals which have the following form:

$$\int_{x_0}^{\infty} dx x^n e^{-x^2} = \int_0^{\infty} dx x^n e^{-x^2} + \int_{x_0}^{0} dx x^n e^{-x^2}$$
$$= \frac{1}{2}\Gamma_f\left(\frac{n+1}{2}\right)[1+(-1)^n] + \frac{(-1)^{n+1}}{2}\Gamma_f\left(\frac{n+1}{2}, \frac{v_o^2}{2\sigma^2}\right), \tag{21}$$

where $\Gamma_f(x_1)$ and $\Gamma_f(x_1, x_2)$ are the standard Gamma functions of one or two arguments [2]. In the limit $\sigma \mapsto 0$ the double argument Gamma function approaches zero exponentially ($\sim e^{-x_0^2}$) since $x_0 \mapsto -\infty$, and hence that contribution can be neglected. Thus, we have:

$$\int_{x_0}^{\infty} dx x^n e^{-x^2} \approx \frac{1}{2}\Gamma_f\left(\frac{n+1}{2}\right)[1+(-1)^n]. \tag{22}$$

Taking this into account, we can write the result for the integral $I$ as

$$I = \sqrt{2\sigma^2}\left(\sqrt{\pi}G(v_o) + \sum_{n=1}^{\infty}\frac{(\sqrt{2}\sigma)^n}{2n!}[1+(-1)^n]\Gamma_f\left(\frac{n+1}{2}\right)G^{(n)}(v_o)\right). \tag{23}$$

Because of the factor $[1+(-1)^n]$, uneven powers of $n$ in the above series disappear.
